# Phosphatases of regenerating liver balance F-actin rearrangements during T cell activation through WD repeat containing protein 1

**DOI:** 10.1101/2023.12.28.573244

**Authors:** Óscar Aguilar-Sopeña, Carlos Carrasco-Padilla, Álvaro Gomez-Moron, Patricia Castro-Sánchez, Matías Estarás-Hermosel, Salvatore Valvo, Sergio Alegre-Gómez, Javier Redondo-Muñoz, Raúl Torres, Sandra Rodríguez-Perales, Shehab Ismail, Francisco Sánchez-Madrid, Noa B. Martín-Cófreces, Michael L. Dustin, Pedro Roda-Navarro

## Abstract

Phosphatases of regenerating liver (PRLs) have been proposed to regulate actin dynamics in lymphoid cells. However, the mechanism mediating this role remained unknown. Here we showed the interaction of the PRLs with the actin regulator WD repeat containing protein 1 (WDR1). The interaction of the PRLs with WDR1 was dependent on F-actin integrity and the proper recruitment of the PRLs to cell membranes through the CAAX motif. Endogenous PRLs and WDR1 were distributed to the Immunological Synapse (IS) and perturbed expression of PRL-1 or PRL-2 by CRISPR/Cas9 mediated genome editing showed that the PRLs were required for proper distribution of WDR1 to filamentous (F)-actin at the IS. Further, perturbed expression of the PRLs or WDR1 resulted in defective IS assembly with deregulated F-actin rearrangements, altered positioning of LFA-1, and reduced IL-2 production. Interestingly, balanced expression of PRLs was required for accumulation of CD3ε at the IS and early activating signalling. We propose that PRLs regulate early T cell signalling and are required for proper F-actin dynamics by regulating the WDR1 access to F-actin networks. As a consequence, PRLs regulate LFA-1 positioning at the IS and proper cytokine secretion.

## Introduction

Activation of T cells involves the formation of a specialized adhesion between T cells and antigen presenting cells (APCs), called the immunological synapse (IS) (Dustin and Choudhuri, 2016). The mature IS is formed by 3 supramolecular activation clusters (SMACs) (Monks et al., 1998): the central SMAC (cSMAC), which is enriched in T Cell Receptor (TCR) and other signalling molecules like protein kinase C (PKC) (Yokosuka et al., 2008) and where signal termination and generation of extracellular vesicles take place (Varma et al., 2006; Vardhana et al., 2010; Choudhuri et al., 2014; Kvalvaag et al., 2023), peripheral SMAC (pSMAC), which is formed by a ring of integrins, such as lymphocyte function-associated antigen 1 (LFA-1) (Cassioli et al., 2021), and dynamic actomyosin arcs (Murugesan et al., 2016), and distal SMAC (dSMAC) characterised by a filamentous (F)-actin retrograde flow, which, together with the actomyosin cytoskeleton at the pSMAC, allows a centripetal movement of activating TCR signalling complexes and integrins towards the cSMAC and pSMAC, respectively (Roy and Burkhardt, 2018). Signalling downstream the TCR and CD28 initiates F-actin rearrangements and the microtubule-organizing centre (MTOC) polarisation to the IS (Martin-Cofreces et al., 2021). This cytoskeleton dynamics is essential for the establishment of a mature IS, which contributes to sustained signalling needed for full T cell activation (Martin-Cofreces et al., 2008), and to intercellular communication based on cytokines, miRNAs or cytotoxic substances (Huse et al., 2006; Stinchcombe et al., 2006; Mittelbrunn et al., 2011; Balint et al., 2020).

Phosphatases of regenerating liver (PRLs: PRL-1, PRL-2 and PRL-3; encoded by *PTP4A1*, *PTP4A2* and *PTP4A3* genes) are dual-specificity phosphatases (DSPs) that belong to the family of protein tyrosine phosphatases (PTPs) (Castro-Sanchez et al., 2019). PRLs have been proposed to regulate the cytoskeleton in cancer cells and during the activation of T cells (Fiordalisi et al., 2006; Nakashima and Lazo, 2010; Castro-Sanchez et al., 2018). Substrates of PRLs in T cells are not characterised and, as members of the family of PTPs, the catalytic activity is thought to occur through an anion thiolate and an aspartic acid residue, which is essential for the release of the product after catalysis (Castro-Sanchez et al., 2019). Proper distribution of PRLs at endo and plasma membranes is mediated by a polybasic region and a C-terminal CAAX motif located downstream to the phosphatase domain (Castro-Sanchez et al., 2020). PRL-2 has been recently proposed to have an important role in T cell development in the thymus (Kobayashi et al., 2017) and our group have described the distribution of PRL-1 and PRL-3 at the IS and a regulatory role of the catalytic activity of the PRLs in actin cytoskeleton and IL-2 production (Castro-Sanchez et al., 2018; Aguilar-Sopena et al., 2020). However, the molecular mechanism mediating the regulation of the actin cytoskeleton by the PRLs has not been established.

Here we have searched for molecular interactions of the PRLs by biochemical approaches and found the interaction of these DSPs with WD repeat containing protein 1 (WDR1), a regulator of the actin turnover by promoting the severing activity of cofilin (Ono, 2018). We have studied the distribution of the PRLs and WDR1 to the IS, the regulation of the subcellular distribution of WDR1 by the PRLs and the regulatory role of these interacting molecules during T cell activation, IS assembly and effector function. We contribute experimental evidences indicating that PRLs balance F-actin rearrangements to tune IS assembly and effector function of T cells via the interaction with WDR1.

## Results

### The PRLs interact with WDR1 at actin filaments

In order to study the molecular mechanism mediating the regulation of F-actin dynamics by PRLs during T cell activation, we initially searched for molecular partners of PRL-1 in Jurkat (JK) cells by a stable isotope labelling using amino acids in cell culture (SILAC) approach. Isotope-labelled cells were transiently transfected with plasmids coding for GFP, PRL-1 fused to the green fluorescent protein (GFP-PRL-1), and a GFP-PRL-1_D72A catalytic death mutant, which is expected to trap substrates (Tonks, 2006). Cells were activated for 5 minutes on plates coated with anti-CD3ε and anti-CD28 monoclonal antibodies along with chimeric ICAM1 in order to trigger TCR, co-stimulation and integrin adhesion. GFP-fusion proteins were precipitated from protein extracts and isotope-labelled peptides of co-precipitating proteins were analysed. Precipitates of the GFP-PRL-1_D72A expressing cells were enriched in peptides of WD-repeat containing protein 1 (WDR1), Coronin 1A and cofilin (**Supplementary Figure 1** and **Supplementary Table 1**). We focused in this finding due to the coordinated function of these three proteins in the regulation of F-actin turnover. The high score found for WDR1 in the GFP-PRL-1_D72A mutant (similar to the score of PRL-1 peptides), prompted us to further proved the interaction of PRL-1 with this protein. GFP-PRL-1, GFP-PRL-1_D72A or the GFP alone were transiently overexpressed in HEK293 cells and GFP precipitates were analysed by Western blot for the presence of WDR1. A mutant of PRL-1 lacking the CAAX motif (GFP-PRL-1_ΔCAAX), which is essential for targeting PRLs to cell membranes and the IS after farnesylation (Castro-Sanchez et al., 2018; Castro-Sanchez et al., 2020), was included to evaluate whether the membrane targeting of PRL-1 was mediating the interaction. Supporting peptide detection in proteomics, WDR1 was detected in precipitates of protein extracts of cells transfected with GFP-PRL-1_D72A and GFP-PRL-1, but not in those extracts of cells transfected with the GFP alone or the GFP-PRL-1_ΔCAAX (**Figure 1A**). These data strongly indicated that the interaction of PRL-1 with WDR1 in HEK293 cells required the targeting of PRL-1 to cell membranes through the farnesylation of the CAAX motif. To further prove the requirement of PRL-1 farnesylation for the interaction, HEK293 cells were treated with the farnesyl transferase inhibitor SCH66336, which hamper the localisation of PRLs to cell membranes (Wang et al., 2002). Due to the high identity among the three PRLs, the interaction of WDR1 with PRL-2 and PRL-3 was also evaluated. Consistent with data obtained with the GFP-PRL-1_ΔCAAX mutant, the interaction of WDR1 and GFP-PRL-1 was substantially reduced in cells treated with SCH66336, indicating that targeting of PRL-1 to cell membranes is required to achieve proper interaction (**Figure 1B**). WDR1 was also detected in precipitates of cells transfected with mCitrine-PRL-2 and GFP-PRL-3 fusion proteins (**Figure 1B**), suggesting that the interaction with WDR1 is a general property of the PRLs.

**Figure 1.**
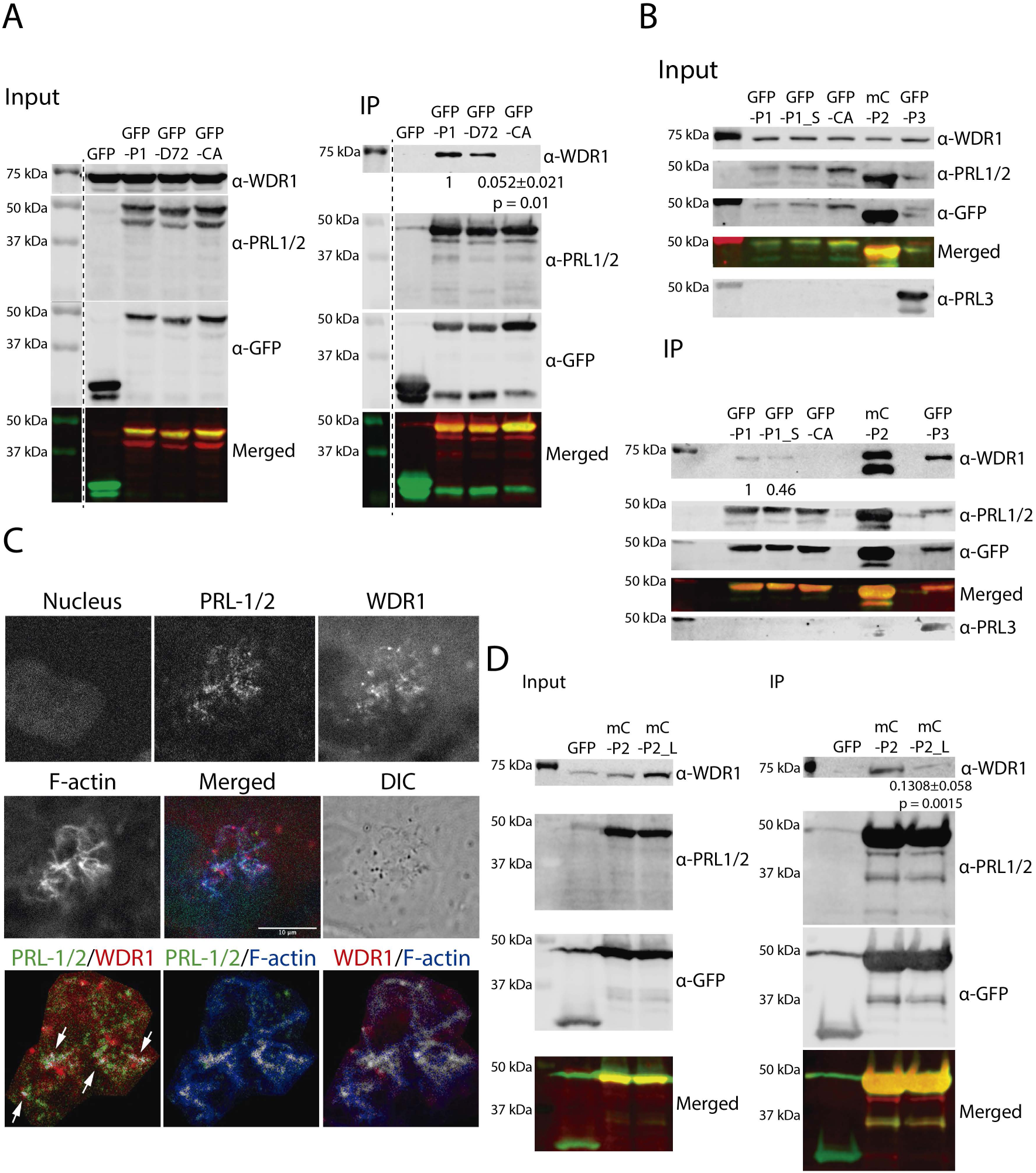
PRLs-WDR1 interaction is dependent on PRLs membrane anchoring and F-actin integrity. **A**. Immunoprecipitation and Western blot of protein extracts of HEK293 cells transfected with GFP, GFP-PRL-1 (GFP-P1), GFP-PRL-1_D72A (GFP-D72) or GFP-PRL-1_ΔCAAX (GFP-CA). The signal of the immunoprecipitated band of GFP-PRL-1_ΔCAAX condition was normalized to the control (GFP-PRL-1) and the mean ± the standard deviation (SD) of n = 4 independent experiments is indicated. Samples (GFP-PRL-1 vs GFP-PRL-1_ΔCAAX) were compared by a one sample t-test (p=0.01). Dashed lines indicate non-showed lanes of the gel. **B**. Immunoprecipitation and Western blot of protein extracts of HEK293 cells transfected with GFP-PRL-1 (GFP-P1) treated or not with SCH (S) (sample GFP-P1_S), GFP-PRL1_ΔCAAX (GFP-CA), mCit-PRL-2 (mC-P2) or GFP-PRL-3 (GFP-P3). Numbers indicate the densitometry of GFP-P1_S normalised to the control. A representative experiment is shown. **C**. Confocal sections of a representative Hela cell (out of n=7 analysed) stained for WDR1 (red channel), F-actin (blue channel) and PRL-1/2 (green channel). The differential interference contrast (DIC) is also shown. Scale bar 10 μm. Lower panels show the pixel maps of the colocalisation of indicated proteins. Arrows point discrete areas of colocalisation between PRL- 1/2 and WDR1. **D**. Immunoprecipitation and Western blot of cell extracts of HEK293 cells transfected with GFP and mCit-PRL-2 (mC-P2) treated or not with latrunculin-A (L) (sample mC-P2_L). The antibody used is indicated. A representative experiment out of n = 3 done is shown. The mean ± standard deviation of the normalised densitometry is shown under the lane of the mC-P2_L. The mC-P2 and mC-P2_L samples were compared by a one sample t-test (p=0.0015).

WDR1 is an F-actin binding protein and then we investigated whether the interaction with PRLs occurred in the context of F-actin. The localisation of endogenous WDR1 and PRLs was studied in Hela cells, which express higher levels of PRL-1 than lymphoid cells (Castro-Sanchez et al., 2018). Cells were co-stained for F-actin and WDR1, along with PRL-1 and PRL-2 by using an antibody specific for both PRLs (see material and methods). Confocal microscopy optical sections placed at the apical site of cells, where F-actin enriched membranes are typically detected, revealed that PRL-1/2 and WDR1 colocalised at discrete points of F-actin (**Figure 1C**; white arrows). To study whether the interaction was dependent on the integrity of F-actin, HEK293 cells overexpressing mCit-PRL-2 were treated with latrunculin-A (Lat-A). The interaction of WDR1 with mCit-PRL-2 was then studied in pull-down experiments as before. Lat-A treatment drastically diminished F-actin levels (**Supplementary Figure 2**) and impaired the interaction of WDR1 with mCit-PRL-2 (**Figure 1D**). Together, our data showed that the interaction of membrane bound PRLs with WDR1 occurred at F-actin and suggested a regulatory role of the interaction in F-actin dynamics.

### PRLs and WDR1 colocalise with F-actin and distribute to the immunological synapse

The interaction of WDR1 and the PRLs was then further studied in JK cells. In order to avoid the perturbation of natural stoichiometry of molecular complexes due to overexpression (Gibson et al., 2013), we took advantage of JK cells expressing endogenous levels of GFP-PRL-1 generated by CRISPR/Cas9 genome editing (so called G-P cells) (Castro-Sanchez et al., 2020). Pull-down experiments were done as before and consistent with an interaction dependent in PRL-1 farnesylation, WDR1 was more strongly detected in GFP-PRL-1 precipitates obtained from control than from SCH66336-treated G-P cells (**Figure 2A**). Then, the distribution of the endogenous PRLs, WDR1 and F-actin was studied by confocal microscopy in resting JK cells and at the IS assembled by these cells and Staphylococcal Enterotoxin E (SEE)-loaded Raji B cells used as APCs. WDR1, PRL-1/2 and F-actin staining showed that both WDR1 and the PRLs colocalised and also overlapped F-actin signal, as estimated by Pearson coefficients (**Figure 2B**, and **2D-2F**). Thus, these data supported the interaction of WDR1 and membrane bound PRLs with F-actin in JK cells. The mature IS in cell-cell conjugates was identified by the peripheral ring of F-actin typically obtained by phalloidin staining. Pearson coefficients indicated that the colocalisation of WDR1 with F-actin increased with the stimulation at any stage of the assembled IS, mature or non-mature. By contrast, the co-localization of PRL-1/2 with either F-actin or WDR1 decreased upon stimulation, reaching the lowest colocalisation at the mature IS (**Figure 2C**-**2F**). Three-dimensional reconstructions of x/y confocal section stacks acquired at cell-cell conjugates clearly located WDR1 at peripheral F-actin of the IS (**Figure 2C**). Conjugates stained for WDR1 and F-actin, along with CD3ε (to localise the cSMAC) or LFA-1 (to localise the pSMAC) further showed the colocalisation of WDR1 at pSMAC and dSMAC areas of the IS, being excluded from the cSMAC (**Figure 2G** and **2H**). These data suggested a regulatory role of the interaction of the PRLs and WDR1 at the F-actin networks organised during T-cell activation and the assembly of the IS.

**Figure 2.**
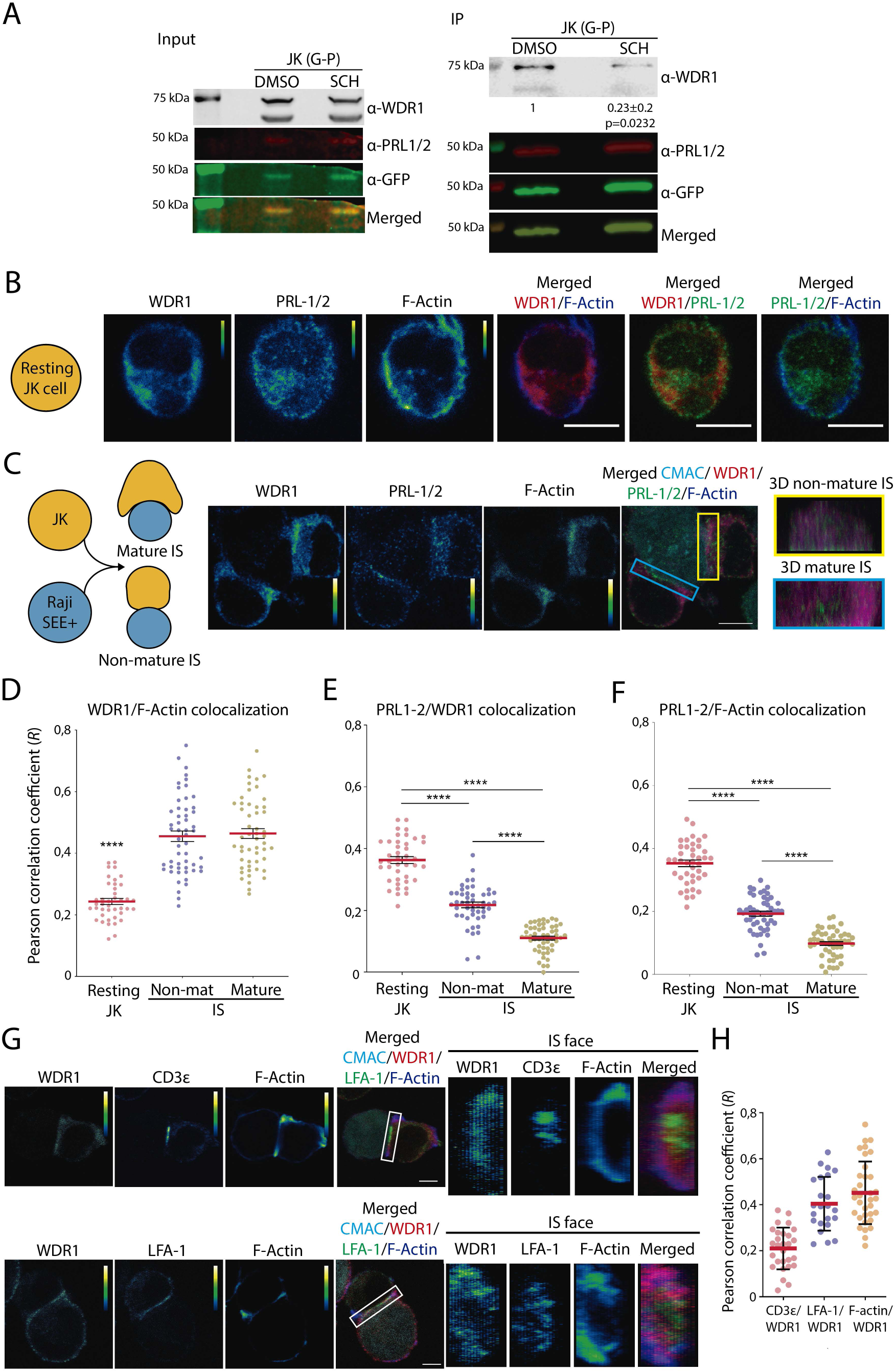
Endogenous PRLs interacted with WDR1 in JK cells and are distributed to the immunological synapse. **A**. Immunoprecipitation of GFP-PRL-1 and Western blot for WDR1 detection in JK G-P cells. Results obtained with protein extracts of cells treated or not with SCH are shown. All the proteins detected by Western blot are indicated. The mean ± standard deviation of the normalised densitometry is shown under the lane of the SCH treated condition. The control (DMSO) and SCH samples were compared by a one sample t-test (p=0.0232). **B** and **C**. Colocalisation of WDR1, PRL-1/2 and F-actin in resting and activated JK cells. Representative resting JK cell (B) and cell conjugate of JK cells interacting with SEE-loaded Raji cell (C). Confocal sections of the CMAC (cyan), red (WDR1), green (PRL-1/2) and blue (actin) channels are shown. Merged and individual (pseudocolor) images are shown. Calibration bar of the pseudocolor is indicated. Scale bar 5 μm. **D, E** and **F**. Graphs indicate the Pearson coefficients for assessing the colocalisation between the different stained molecules. Dots indicate values of individual synapses or resting cells obtained from n = 2 experiments. Means ± standard errors are shown. Samples were compared by a one-way ANOVA with a Tukey’s multiple comparison test. ****p<0.001. **G**. Representative cell conjugates of JK cells activated by SEE-presentation. The red (WDR1), green (CD3ε or LFA-1) and blue (F-actin) channels are shown in both as pseudocolor and as a merged image, also at the IS face. Calibration bar of the pseudocolor is indicated. Scale bar 5 μm. **H**. Graphs indicate the Pearson coefficients for assessing the colocalisation between CD3ε, LFA-1 or F-actin and WDR1. Dots indicate values of individual synapses obtained from n = 2 experiments. Means ± standard deviations are shown.

### The PRLs regulate WDR1 distribution to F-actin networks

To study the possible regulation of WDR1 by PRL-1 and PRL-2, knockout cells for *PTP4A1* and *PTP4A2* genes were generated by CRISPR/Cas9-mediated genome editing in the JK cell line. Interestingly, the analysis of PRL-1 and PRL-2 expression by Western blot revealed a compensated expression of both proteins in initial edited cells, knockout clones and a pool of clones established to avoid clonal effects (**Figure 3A** and **Supplementary Figure 3A-3C**). This suggested a shared regulatory mechanism of these proteins for the viability of JK cells.

**Figure 3.**
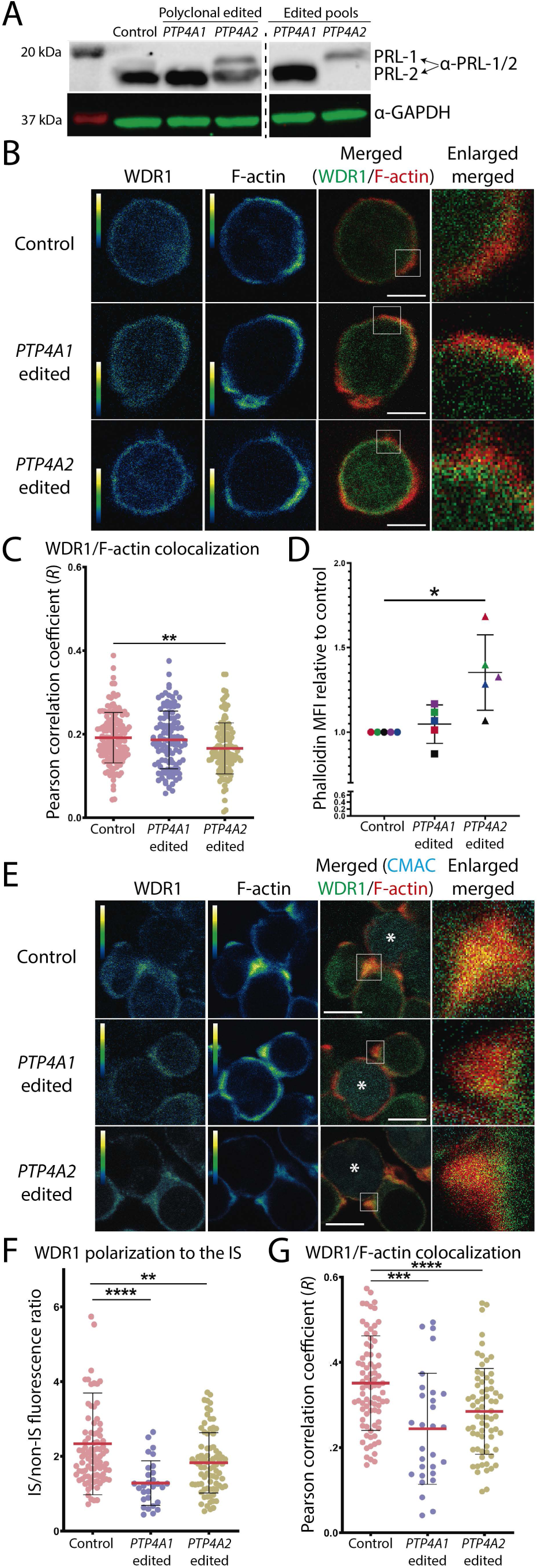
PRLs are necessary for a proper distribution of WDR1 to the F-actin. A. Expression of endogenous PRL-1/2 assessed by Western blot of non-transfected JK control cells, *PTP4A1* and *PTP4A2* edited polyclonal populations and *PTP4A1* and *PTP4A2* knockout pools. Molecular weights and detected proteins are indicated. **B.** Confocal sections of representative control, *PTP4A1* and *PTP4A2* edited JK cells. The red (WDR1) and green (F-actin) channels (pseudocolor), as well as the merged image are shown. A zoomed area (white square) of the merged image is shown. Calibration bar of the pseudocolor is indicated. Scale bar 5 μm. **C**. Graphs indicating the Pearson coefficients to assess the colocalisation between WDR1 and F-actin in each sample at the IS. Dots indicate individual cells analysed and the mean ± standard deviation from n=2 independent experiments is shown. Samples were compared by a one-way ANOVA with a Tukey’s multiple comparison test. **p<0.01. **D**. F-actin levels assessed by Phalloidin staining of each sample by flow cytometry. Coloured symbols indicate the n=5 independent experiments done. Data were normalized to the control and compared by a one sample t-test. *p<0.05. **E**. Confocal sections of representative cell conjugates of JK cells interacting with SEE-loaded and Raji cells. The red (WDR1) and green (F-actin) channels (pseudocolor), as well as the merged images are shown. A zoomed area (white square) of the merged image is shown. Calibration bar of the pseudocolor is indicated. Scale bar 5 μm. **F**. WDR1 polarisation to the IS. **G**. Pearson coefficients to assess the colocalisation between WDR1 and F-actin in each sample at the IS. In F and G, dots indicate individual cell conjugates obtained from n=3 independent experiments and the mean ± standard deviation is shown. Samples were compared by a one-way ANOVA with a Tukey’s multiple comparison test. **p<0.01, ***p<0.001 and ****p<0.0001.

Then, we studied the distribution of WDR1 to F-actin in *PTP4A1* or *PTP4A2* edited cells. Confocal microscopy and Pearson coefficients showed a decreased colocalisation of WDR1 with F-actin in *PTP4A2* edited cells in comparison with what found in *PTP4A1* edited or control cells (**Figure 3B** and **3C**). Due to a deficient recruitment of WDR1 to F-actin in *PTP4A2* edited cells, it was expected deficient activity of cofilin and increased amount of F-actin in these cells, and this was found by F-actin staining and flow cytometry (**Figure 3D**). Thus, PRL-2 seemed to control actin homeostasis in non-stimulated JK cells by assisting the recruitment of WDR1 to F-actin sites.

We further studied the role of the PRLs in the recruitment of WDR1 to the IS, where they are expected to regulate F-actin networks. JK cells were stimulated with SEE-loaded Raji B cells as before and the polarisation of WDR1 and F-actin to the IS in edited or control cells was evaluated. Colocalisation of WDR1 and F-actin was found at peripheral areas of the IS established by control cells. However, a reduced polarisation of WDR1 to the IS and colocalisation with F-actin was found in both *PTP4A1* and *PTP4A2* edited cells when compared to control cells (**Figure 3E-3G**). These data indicated that both PRLs are required for the correct local recruitment of WDR1 to F-actin networks at the IS.

### PRLs and WDR1 regulate F-actin rearrangements at the IS

The regulatory role of PRL-1, PRL-2 and WDR1 in the F-actin rearrangements at the IS was then investigated. To this aim, we also perturbed the expression of WDR1 to directly evaluate the role of WDR1 in this process. CRISPR/Cas9-mediated genome editing of JK cells was done as before. *WDR1* edited JK cells showed a marked reduced expression of WDR1 as assessed by Western blot and several mutations of the protein were detected by DNA sequencing (**Supplementary Figure 4A and 4B**). To study the F-actin rearrangements at the mature IS in *PTP4A1*, *PTP4A2* and *WDR1* edited cells, T-cell activation was evaluated in T cell-APC conjugates under confocal microscopy as before. *PTP4A2* and *WDR1* edited cells showed an increased polymerization of F-actin at the IS of cell-cell conjugates (**Figure 4A** and **4B**). This is consistent with a requirement of PRL-2 and WDR1 to achieve proper F-actin severing and depolymerisation during T cell stimulation. By contrast, this was not observed in *PTP4A1* edited cells, in which a slight reduction of F-actin polymerization was found in comparison to control cells (**Figure 4A** and **4B**), suggesting an additional regulatory role of the PRLs during TCR cell stimulation. Remarkably, the polarisation of F-actin in *PTP4A1* and *PTP4A2,* but not *WDR1*, edited cells paralleled an altered accumulation of CD3ε at the IS (**Figure 4A** and **4C**). Thus, balanced expression of PRL-1 and PRL-2 seems to control CD3ε distribution at cognate cell-cell interactions.

**Figure 4.**
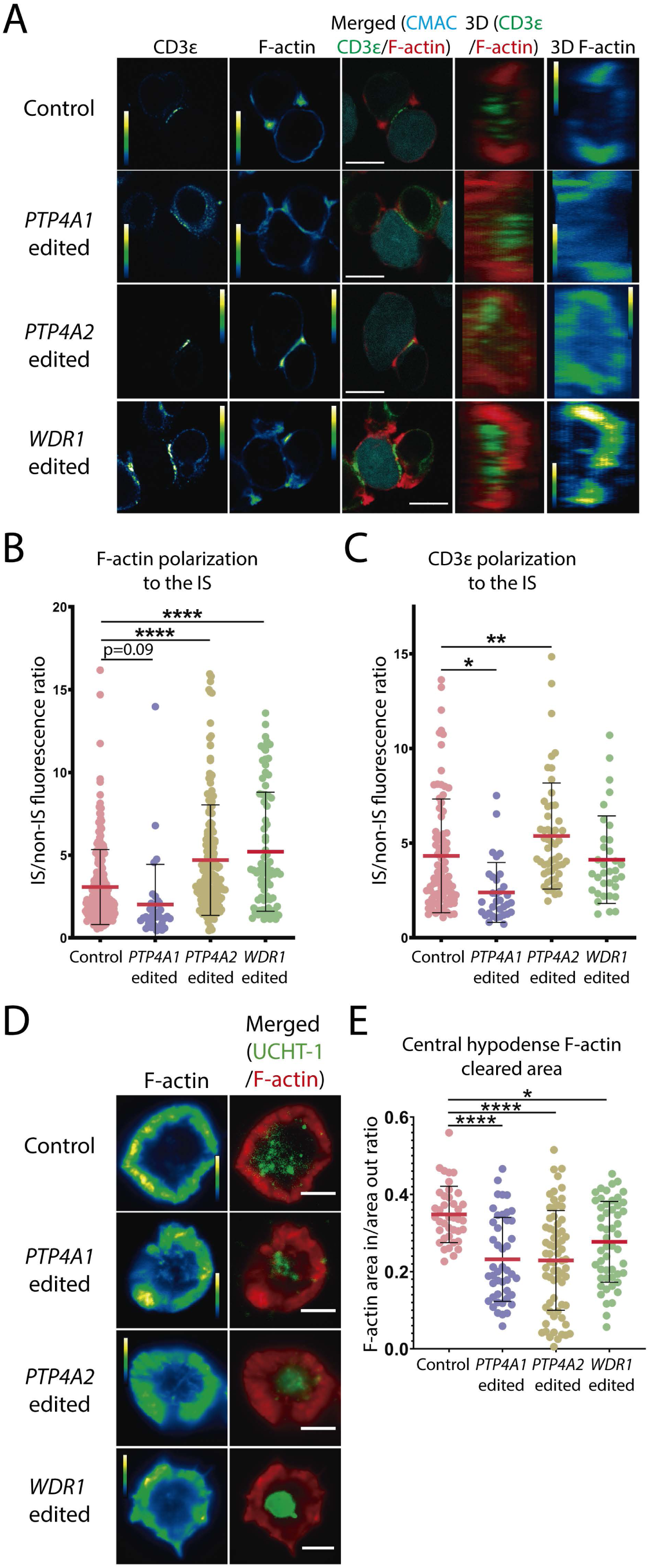
PRLs and WDR1 regulate actin rearrangements at the IS. **A**. Confocal sections of representative conjugates of JK (control or *PTP4A1*, *PTP4A2* or *WDR1* edited) and SEE-Raji cells IS. The green (CD3ε) and red (F-actin) channels in pseudocolor and the merged image including the cyan channel (CMAC) are shown. The calibration bar of the pseudocolor is indicated. Scale bar corresponds to 5 μm. 3D reconstructions of the merged image and pseudocolor F-actin channel are shown. **B**. F-actin polarisation calculated from confocal sections. **C**. CD3ε polarisation calculated from confocal sections. In B and C, dots indicate individual conjugates analysed and the mean ± standard deviation in each sample from at least n=2 independent experiments are shown. Samples were compared by a one-way ANOVA with a Tukey’s multiple comparison test. *p<0.05, **p<0.01 and ****p<0.0001. P values marginally significant are also indicated. **D**. TIRF microscopy images of representative JK cells (control or *PTP4A1*, *PTP4A2* or *WDR1* edited) activate on ALBs. F-actin labelling (pseudocolor) and merged images with UCHT-1 (green) and F-actin (red) is shown. Pseudocolor and scale bar are shown. **E**. Central hypodense F-actin cleared area of each analysed interaction. Dots represent analysed cells and it is indicated the mean ± standard deviation in each sample from n=3 independent experiments. *p<0.05 and ****p<0.0001. Samples were compared by a one-way ANOVA with a Tukey’s multiple comparison test.

Due to the expected role of WDR1 in F-actin severing and depolymerisation, the establishment of the hypodense F-actin central area of the IS was evaluated. In this case, T cells were activated on activating lipid bilayers (ALBs) containing mobile fluorescently-labelled anti-CD3ε and chimeric ICAM-1 to define de mature IS. ALBs avoided the presence of the APC F-actin, achieving a more reliable assay. The assembled IS was tracked by Total Internal Reflexion Fluorescence microscopy (TIRFm) to restrict chromophore excitation to the activation site. In these experiments, any of the edited cells could established a completely open hypodense F-actin network at the central areas of the IS (**Figure 4D** and **4E**), indicating a perturbed actin dynamics at the mature IS. To extend our study to primary cells being activated on ALBs, F-actin networks were evaluated in peripheral blood CD4 T cells with reduced expression of PRL-1, PRL-2 and WDR1 after mutations introduced at the encoding genes following the same CRISPR/Cas9 genome editing strategy (**Supplementary Figure 3D** and **Supplementary Figure 4C**). WDR1 localisation at F-actin areas was also evaluated in *PTP4A1* and *PTP4A2* edited cells. Consistent with data obtained in JK cells, CD4 T cells with *PTP4A1* and *PTP4A2* edited genes showed a reduced colocalisation of WDR1 with F-actin (**Figure 5A and B**). Moreover, any of the edited cells established completely open hypodense F-actin areas at the activating surface of the cell (**Figure 5A** and **5C**). These data indicated that PRLs are required for a proper organisation of hypodense F-actin networks at the central area of the IS by assisting the distribution of WDR1 to F-actin sites. Together, our data showed that the proper expression of WDR1 is required for proper F-actin rearrangements at the mature IS and that PRLs regulate this process by controlling the access of WDR1 to F-actin sites.

**Figure 5.**
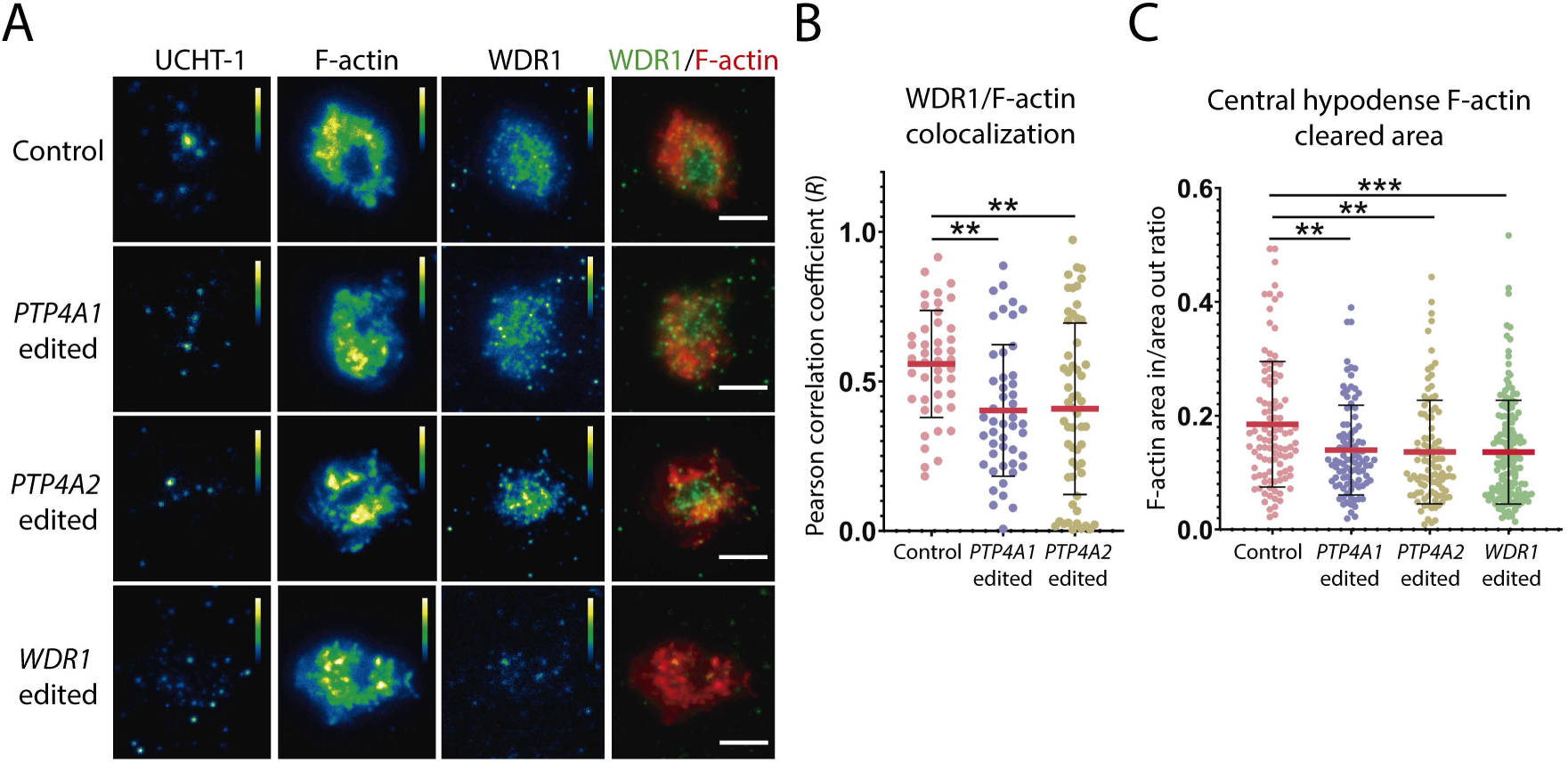
PRLs and WDR1 regulate F-actin rearrangements at the IS in primary cells. **A**. TIRF microscopy images of representative CD4+ T cells (control or edited to knockout PRL-1, PRL-2 or WDR1 encoding genes) activated on ALBs. The UCHT-1, F-actin and WDR1 channels are displayed in pseudocolor. Pseudocolor bar is shown. The green WDR1 and red F-actin merged are shown. Scale bar corresponds to 5 μm. **B**. Graph indicates Pearson coefficients to assess the colocalisation between WDR1 and F-actin in each condition. **C**. Graph indicates the central hypodense F-actin cleared area of each analysed interaction. In B and C, dots represent individual cells analysed, the mean ± standard deviation of each sample from n=2 independent experiments are shown, and samples were compared by a one-way ANOVA with a Tukey’s multiple comparison test. **p<0.01 and ***p<0.001.

### The PRLs and WDR1 regulates LFA-1 distribution at the IS

Due to the distribution of WDR1 to the peripheral areas of the IS (**Figure 2F**), the perturbed F-actin networks found at the mature IS in edited cells and the essential role of F-actin networks in LFA-1 positioning (Comrie et al., 2015), we evaluated the localization of this integrin in control, *PTP4A1*, *PTP4A2* or *WDR1* edited cells stimulated by SEE-loaded Raji cells. Confocal microscopy showed a discrete colocalisation of LFA-1 with internal sites of the F-actin ring at the dSMAC in wild-type JK cells, indicating the well-known location of this integrin at the pSMAC. Interestingly, all edited cells showed an increased localisation of LFA-1 at distal F-actin as indicated by a higher number of LFA-1 particles at distal F-actin (see material and methods) and higher Pearson coefficients to estimate colocalisation (**Figure 6A-6C**). These data suggested an essential role of the PRLs and WDR1 in the proper positioning of LFA-1 at the pSMAC. We then further investigated the integrin positioning at the IS by measuring the distribution of the LFA-1 ligand ICAM-1 at ALBs under TIRFm. Consistent with a hampered location of LFA-1 at the pSMAC, the area of recruited ICAM-1 found under *PTP4A1*, *PTP4A2* or *WDR1* edited cells was higher than the one observed under control cells (**Figure 6D** and **6E**). These data indicated that PRLs and WDR1 control proper actin cytoskeleton rearrangements allowing the correct positioning of LFA-1.

**Figure 6.**
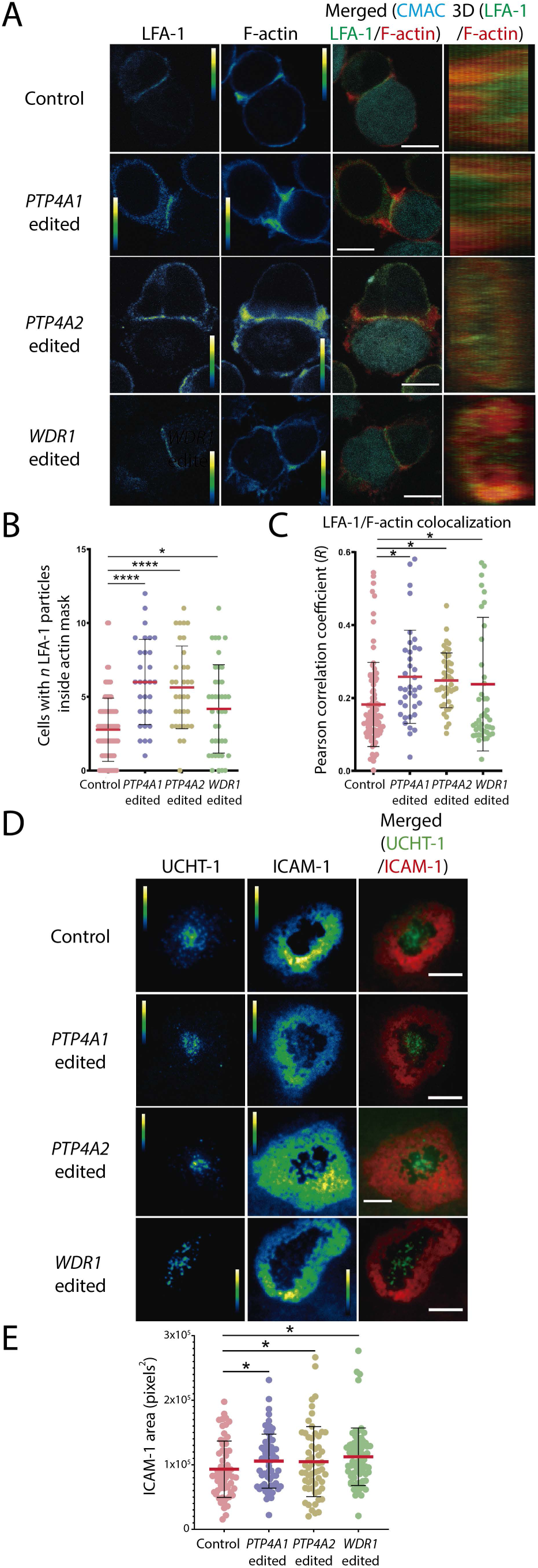
PRLs and WDR1 control LFA-1 distribution to T cell IS. A. Confocal sections of representative conjugates of JK cells (control or *PTP4A1*, *PTP4A2* or *WDR1* edited cells) and SEE-loaded Raji cells IS. The green (LFA-1) and the red (F-actin) channels in pseudocolor, as well as the merged image with the cyan (CMAC) channel are shown. The calibration bar of pseudocolor is indicated. Scale bar corresponds to 5 μm. Synapse surface from 3D reconstructions of confocal sections are displayed with LFA-1 and F-actin channels merged. **B**. Polarisation of LFA-1 to the IS. **C**. Number synapses with *n* LFA-1 particles inside a F-actin mask obtained from the IS surface. In B and C, symbols represent individual cell conjugates analysed. The mean ± standard deviation of each sample from at least n=2 independent experiments are shown. Samples were compared by a one-way ANOVA with a Tukey’s multiple comparison test. *p<0.05 and ****p<0.0001. **D**. TIRF microscopy images of JK cells (control or *PTP4A1*, *PTP4A2* or *WDR1* edited) activated on ALBs. The green (UCHT-1) and red (ICAM-1) channels in pseudocolor and the merged imaged are shown. Pseudocolor calibration bar is shown. Scale bar corresponds to 5 μm. **E**. Area in pixels^2^ of the accumulated ICAM-1 at ALBs. The mean ± standard deviation of each sample from n=2 independent experiments are shown. Samples were compared by a one-way ANOVA with a Tukey’s multiple comparison test. *p<0.05.

### PRLs and WDR1 regulates TCR signalling and cytokine production

Due to altered distribution of CD3ε and F-actin rearrangements found, we aimed to investigate the T cell activation and IL-2 production in *PTP4A1*, *PTP4A2* and *WDR1* edited cells. Interestingly, *WDR1* and *PRL-2* edited cells showed an increased surface levels of CD3ε, CD4, CD28 and LFA-1, indicating a regulatory role in the surface expression of these proteins. The increased expression of CD28 and LFA-1 was also observed in *PTP4A1* edited cells. By contrast increased surface levels of CD3ε and CD4 were not observed; even CD3ε expression levels seemed to be lower in these edited cells (**Supplementary Figure 5**).

Early signalling upon activation was evaluated via ERK1/2 activation assessed by Western blot. A general reduced ERK1/2 signalling was found in *PTP4A1* edited cells; while it was enhanced at early times in *PTP4A2* and *WDR1* edited cells (**Figure 7A**). Activation of control and edited JK cells was evaluated by CD69 surface staining and flow cytometry after stimulation with growing amounts of SEE-loaded in Raji cells. The SEE-induced surface expression of CD69 was higher in *WDR1* and *PTP4A2* edited cells than in control and *PTP4A1* edited cells (**Figure 7B**). Thus, these data indicated that the PRLs tune early events of T cell activation, which also seems to required adequate levels of WDR1. We have previously proposed that the catalytic activity of the PRLs is required for a proper production of IL-2 (Castro-Sanchez et al., 2018; Aguilar-Sopena et al., 2020). We then studied by ELISA the amount of secreted IL-2 in cell supernatants as a consequence of the stimulation. Interestingly, although the T cell stimulation was enhanced in *PTP4A2* and *WDR1* edited cells, the production of IL-2 was reduced in all *PTP4A1*, *PTP4A2* and *WDR1* edited cells (**Figure 7C**). Therefore, while PRL-2 and WDR1 seemed to control initial TCR SEE-triggered signalling and CD69 induced surface expression, PRL-1 seems to be necessary for proper early signalling. Interestingly, the PRLs and WDR1 seemed to be required for a proper production of IL-2 measured in cell supernatants, indicating a common function in later events during T cell effector function.

**Figure 7.**
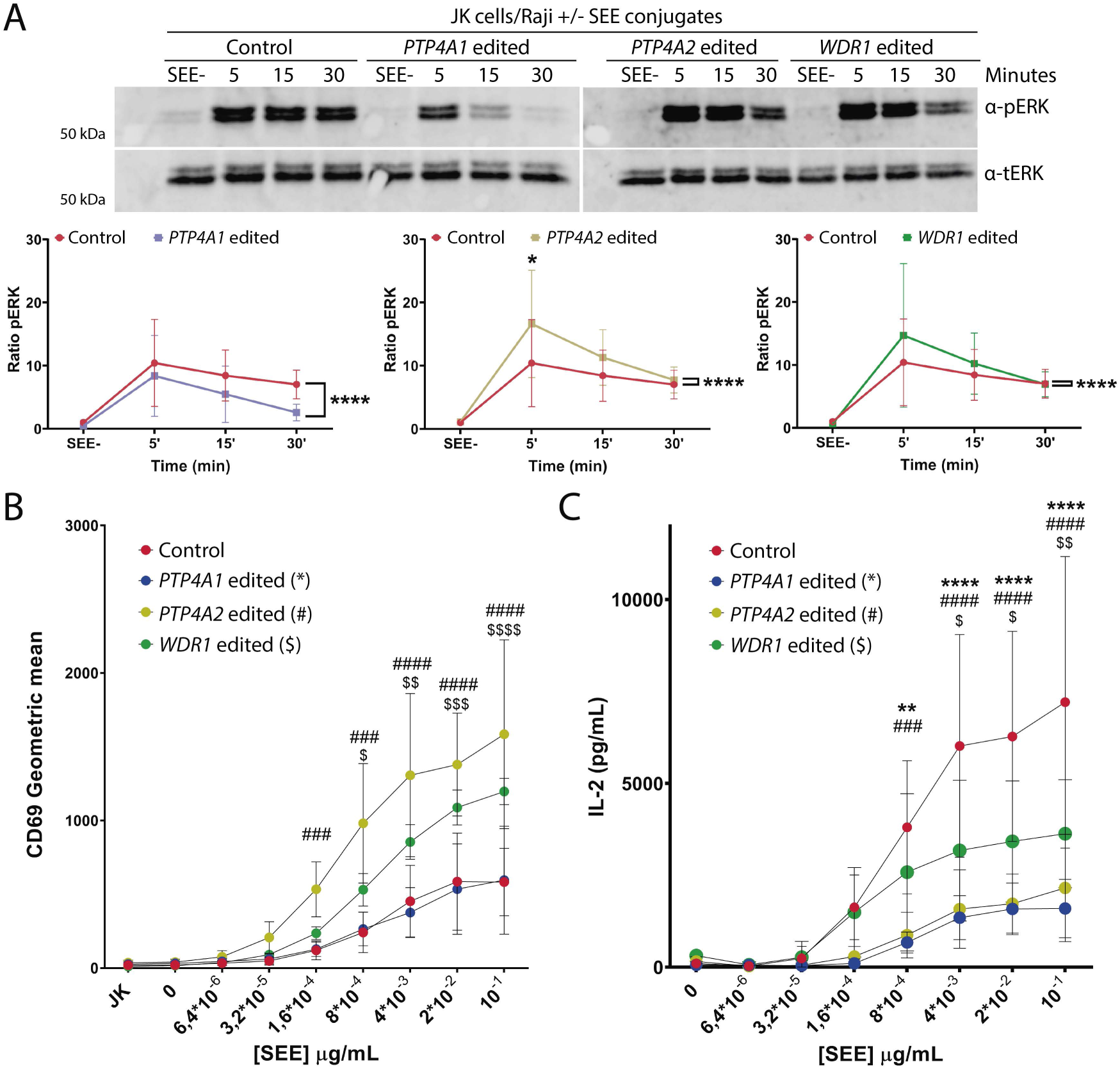
PRLs and WDR1 contributes to T cell activation. **A**. Western blots showing the time course of phosphorylation of pT202/Y204 ERK1/2 in control or *PTP4A1*, *PTP4A2* or *WDR1* edited cells conjugated with SEE-preloaded or not Raji B cells for the indicated times. Graphs show the mean ± standard deviation of the ratio of phosphorylation to total protein normalised to unstimulated control. Data coming from n=6 independent experiments was compared by a two-way ANOVA. * p<0.05 p and **** p<0.0001. **B** and **C.** CD69 induction and IL-2 secretion analysed by flow cytometry and ELISA, respectively, of the indicated samples of JK cells stimulated for 24 hours with Raji cells loaded with the indicated concentrations of SEE. Graphs indicate the mean ± standard deviation of each sample from n=3 independent experiments. Samples were compared by a one-way ANOVA with a Tukey’s multiple comparison test. The * symbol represents the comparison between *PTP4A1* edited and control cells, the # symbol indicates the comparison between *PTP4A2* edited and control cells and the $ symbol represents the comparison between *WDR1* edited and control cells. */#/$ p<0.05, **/##/$$ p<0.01, ***/###/$$$ p<0.001 and ****/####/$$$$ p<0.0001.

## Discussion

In this work, we provide experimental evidence showing the interaction of phosphatases of regenerating liver with the cytoskeleton regulator WDR1. The PRLs and WDR1 distribute to the IS, where they regulate actin rearrangements and LFA-1 positioning. PRLs and WDR1 are required for proper intracellular signalling and IL-2 production. Our data show a molecular mechanism mediating the regulation of the actin cytoskeleton by the PRLs and WDR1 in lymphoid cells.

By using proteomics and PRL-1 fluorescent fusion proteins, we have identified the interaction of the PRLs with WDR1. We show that the interaction with WDR1 is a general property of the PRLs. Our data indicate that the interaction with WDR1 depends on cell membranes localisation and does not depend on the phosphatase activity. It should be studied whether this interaction depends on the described ability of PRLs to trimerize (Jeong et al., 2005) or whether these proteins can form heterotrimers to achieve certain regulatory roles in the cell. It has been shown that WDR1 interacts with cofilin-decorated actin filaments and enhances filament disassembly (Ono, 2018). Using multi-wavelength single molecule fluorescence imaging, recent studies have observed that WDR1, cofilin and Coronin 1B work through a temporally ordered pathway to induce efficient severing and disassembly of actin filaments. In this model, Coronin 1B binds to actin filaments first, and increases the subsequent binding of cofilin. Then, WDR1 is recruited to cofilin-decorated actin filaments to promote the action of cofilin (Jansen et al., 2015). The interaction of WDR1 and PRLs at discrete sites of F-actin, the decreased location of WDR1 in F-actin and the incremented F-actin observed in *PTP4A2* edited cells indicate that PRLs might contribute a new regulatory layer to tune cofilin function by assisting the access of WDR1 to actin filaments, rendering proper F-actin homeostasis. It is not clear at this point whether the interaction with WDR1 is direct or indirect in the context of the molecular complex regulating F-actin and whether WDR1 or other actin regulators are substrates of the PRLs. Interestingly, it has been shown that WDR1 can be phosphorylated by Src family kinases and dephosphorylated by the protein phosphatase EYA3. In this regard, lack of Src family kinases-mediated phosphorylation of WDR1 induces malformed actin fibbers and F-actin decreased levels (Mentel et al., 2018). Thus, it can be speculated that the PRLs could regulate the activity of Src family kinases in T cells (such as Lck) or directly the phosphorylation state of WDR1, and the missing of these DSPs might produce an increased phosphorylation of WDR1 rendering higher levels of F-actin. However, it is either not known whether phosphorylation events in WDR1 control its function towards cofilin. The regulation exerted by the PRLs might be also independent of the phosphatase activity, like, for example, in the previously reported interaction of the PRLs with CNNM proteins (Gimenez-Mascarell et al., 2017). The mechanism proposed in this work for actin regulation is compatible with this possibility.

The colocalisation of the PRLs and WDR1 is better detected in non-stimulated JK cells and at non-mature cognate interactions with APCs. This suggest that, during T cell activation, there is an early WDR1 regulation by PRLs, which have been proposed to redistribute to the IS during nascent cognate synapses (Castro-Sanchez et al., 2018). At nascent cognate interactions, the PRLs might allow the access of WDR1 to trigger local actin rearrangements required for proper TCR triggering (Wipa et al., 2020), which is needed for the assembly of the mature IS. At the mature IS, the PRLs and WDR1 localise at different zones when confocal microscopy is used, finding only a partial overlapping of these molecules. However, further characterisation of the spatial and temporal regulation of WDR1/PRLs interaction will be needed to deeply understand where and when this interaction takes place. In this regard, it will be important to study how the molecular dynamics at the IS is regulated by these proteins; specially taking into account the previously observed localisation of GFP-fused PRLs at TCR-containing vesicles and LFA-1 areas of the mature IS (Castro-Sanchez et al., 2018; Aguilar-Sopena et al., 2020). Location of WDR1 at pSMAC and internal dSMAC areas of the mature IS suggest a regulatory role of WDR1 in actomyosin arcs and/or distal actin retrograde flow, which are essential for the dynamics of signalling microclusters and positioning of LFA-1 at the pSMAC. Consistent with the localisation of WDR1 and overexpressed PRL-1 at LFA-1 areas of the IS, decreased expression of PRL-1, PRL-2 and WDR1 in genome-edited cells perturbs the positioning of this integrin at the pSMAC. Thus, we propose that the PRLs and WDR1 regulate the F-actin dynamics required for proper LFA-1 positioning at the pSMAC of the IS.

Our data indicate that PRLs and WDR1 regulates early signalling triggered by antigen stimulation. While PRL-2 and WDR1 seem to control T cell activation, PRL-1 seems to be required to achieve adequate levels of ERK1/2 activation. Interestingly, our data indicate that balanced levels of the PRLs might control the traffic of the TCR to the IS and early signalling. Thus, the role of the PRLs on the endosomal compartment should be further evaluated. In addition to a regulatory role on the endosomal compartment, increased signalling observed in *PTP4A2* or *WDR1* edited cells might be a consequence of altered actin dynamics at the IS or caused by an altered secretory pathway in resting cells. Increased expression of surface receptors observed here support this idea. Interestingly, however, all the studied molecules seem to be required for a proper production of IL-2. Then, it is possible that these proteins are required for latter transcription of the *IL-2* gene or IL-2 translation or secretion. Due to the hampered establishment of the hypodense F-actin network found in edited cells, our data suggest a regulatory role of the PRLs and WDR1 in cytokine secretion at the IS, where the IL-2 has been proposed to be secreted (Huse et al., 2006). Interestingly, WDR1 has been proposed to regulate constitutive secretion in non-lymphoid cells (Lopez-Coral et al., 2018) and in our model PRLs might regulate secretion via WDR1. Nonetheless PRL-3 has been proposed to dephosphorylate phosphoinositide PI(4,5)P2 (McParland et al., 2011), which might support adequate PIP3 local levels for F-actin accumulation at the IS (Le Floc’h et al., 2013). A mechanism mediated by PIs might be also accounting for the observed F-actin, IL-2 or LFA-1 phenotype found here in PRL-1 or PRL-2 depleted cells. The regulatory role of PRLs and WDR1 in secretion of lytic granules of cytotoxic lymphocytes should be also investigated. This might uncover new strategies to generate anti-tumour specific CTLs in adoptive transfer strategies (Sadelain et al., 2003).

Mutations of WDR1 have been associated with the development of a primary immunodeficiency (Kuhns et al., 2016; Pfajfer et al., 2018). Consistent with data obtained in edited cells, it has been observed that WDR1 missense mutations cause aberrant actin dynamics and function of both T- and B-cells (Pfajfer et al., 2018). Zebrafish WDR1-deficient neutrophils show hampered nucleus integrity, which is interestingly recover upon Coronin 1A depletion, indicating that the phenotype is associated to an excess of non-functional cofilin at the actin filament (Bowes et al., 2019). WDR1 is also required for a proper chemotactic migration and cytokinesis of neutrophils and Jurkat T cells (Kato et al., 2008). It will be interesting to study the the role of the PRLs in these processes.

In summary, we propose that the interaction of WDR1 and the PRLs constitutes a new axis for the regulation of actin cytoskeleton in T cells. The interaction of PRLs and WDR1 should be further investigated in other immune cell types to better understand the regulatory role of this interaction in the immune system.

## Materials and methods

### Cells

CD4 T cell line Jurkat (JK, clone E6-1, code TIB-152 in atcc.org) was cultured in RPMI 1640 with fetal bovine serum (FBS) (10 %), penicillin (100 U/ml), streptomycin (100 μg/ml), L-Glutamine (2 mM), sodium pyruvate (2 mM) and non-essential aminoacids (1x) (JK medium). Raji B cell line (code CCL-86 in atcc.org) was cultured in RPMI 1640 supplemented with FBS (10 %), penicillin (100 U/ml), streptomycin (100 μg/ml) and L-Glutamine (2 mM). HEK-293 (human embryonic kidney) epithelial cell line (code CRL-1573 in atcc.org) and HeLa (human cervical adenocarcinoma, code in CCL-2 atcc.org) cell lines were cultured in Dulbecco’s modified Eagle’s medium (DMEM) supplemented with FBS (10 %), penicillin (100 U/ml), streptomycin (100 μg/ml) and L-Glutamine (2 mM). All the cell culture reagents were purchased from Lonza (USA).

Peripheral blood of healthy donors was obtained from buffy coats from the John Radcliffe Hospital transfusion centre (Oxford, United Kingdom). For purification of peripheral blood mononuclear cells (PBMCs), blood was diluted two times on phosphate-buffered saline (PBS, Sigma Aldrich, USA) and centrifuged at 600 g, 35 minutes without acceleration or deceleration on lymphocyte isolation solution (Thermofisher, USA). Cells were then washed three times with PBS, resuspended on RPMI 1640 with FBS (10 %), penicillin (100 U/ml), streptomycin (100 μg/ml) and L-Glutamine (2 mM). CD4 T cells were isolated from these PBMCs by negative selection, using the Dynabeads Untouched Human CD4 T cell kits (Thermofisher, USA). T lymphoblast were generated by culturing T cells at 2 million/mL onto P12 wells on RPMI 1640 with FBS (10 %), penicillin (100 U/ml), streptomycin (100 μg/ml) and L-Glutamine (2 mM), in presence of Dynabeads Human T-Activator CD3/CD28 (ratio 1:2, from Thermofisher, USA) and IL-2 (50 U/mL, from Proteintech, Germany) for 4 days. After this time, magnetic Dynabeads were removed with a magnet and 50 U/mL of IL-2 were added every 48 hours, and they were left growing for 6 days.

### Antibodies and reagents

Mouse α-CD3-APC, CD4-PE, CD28-APC and CD69-APC were from BD Biosciences (USA). Mouse CD11a-APC and CD19-PE were from Biolegend (USA). Mouse α-PRL-1/2, α-tubulin and mouse IgG MOPC were from Sigma Aldrich (USA). Mouse α-WDR1 was from Santa Cruz (USA). Rabbit α-WDR1 was from Abcam (UK). Secondary antibodies goat anti-rabbit-Ig FITC, donkey anti-mouse Alexa 488 and goat anti-rabbit Alexa 594 were acquired from Life technologies (USA). All the labelled mouse IgG1 isotype were from Immunostep (Spain). Anti-CD3ε (T3b) and anti-LFA-1 (TP1/40) supernatants were produced at Dr. Francisco Sanchez-Madrid laboratory (Hospital Universitario de la Princesa, Madrid, Spain). α-phospho-p44/p42 MAPK (ERK1/2) and α-total p44/p42 MAPK (ERK1/2) were from Cell Signaling (USA).

### Proteomics

Stable Isotopes Labelling with Amino Acids in Cell Culture (SILAC) was done for mass spectrometric-based quantitative proteomics. Jurkat clone J77 cells were cultured in different mediums (to then been able to differentiate between each condition) which contain different isotopes of the amino acids lysine and arginine (L-Lys 1H (Lys0) and L-Arg 12C6,15N2 (Arg0) for “light medium”; L-Lys 2H4 (Lys4) and L-Arg 13C6,14N4 (Arg6) for “medium medium”; L-Lys 13C6,15N2 (Lys8) and L-Arg 13C6,15N4 (Arg10) for “heavy medium”) several days. Cells cultured in light, medium and heavy medium were transfected with plasmids which encoded the proteins GFP-PRL-1_D72A, GFP and GFP-PRL-1, respectively. The transfection was carried out by nucleofection using Amaxa® Cell Line Nucleofector Kit V (Program X-001, Lonza, USA). At 24 h after transfection, J77 cells were stimulated with surface-coated anti-CD3 and anti-CD28 antibodies and ICAM-1 for 5 minutes. Then, cells were lysed for 30 minutes in ice-cold RIPA buffer [20 mM Tris-HCl pH 7.5; 1% NP-40; 0.5% sodium deoxycholate; 0.1% SDS; 150 mM NaCl; 10 mM β-glicerophosphate; 1X protease inhibitor cocktail; 10 mM NaF; 1 mM PMSF; 1 mM Na3VO4]. The lysates were subjected to immunoprecipitation using GFP-Trap (Chromotek, Germany) by following the company protocol. Next, the resultant pull downs were digested by trypsin to produce a mixture of peptides for Liquid Chromatography–Mass Spectrometry (LC-MS) analysis. The data was done using MaxQuant® software which identifies the proteins whose peptides were found in LC-MS. For each protein, the ratios between the protein found in J77 cells transfected with either GFP-PRL-1 or GFP-PRL-1_D72A and the intensity of this peptides in J77 cells transfected with GFP were calculated.

### Immunoprecipitation and western blot

For pull down experiments, HEK293 cells were transfected using lipofectamine 2000 (Thermofisher, USA) following the manufacturer protocol. Cells were seeded in P6 wells and transfected with the indicated plasmid when the cell confluence was between 70-90%. Before the co-immunoprecipitation assays, transfected cells were cultured for 16 hours and collected with a scraper. Transfection was checked by flow cytometry and cells were lysed in ice-cold lysis buffer (NaCl 150 mM, Tris-HCl pH 7.5 20 mM, NP40 1 %, Triton X100 0.2 %, glycerol-phosphate 5 mM, EDTA 2 mM, MgCl2 1.5 mM and protease inhibitor cocktail [NaF 10 mM, PMSF 1 mM and Na3VO4 2 mM]) for 30 minutes and centrifuged for 10 minutes at 13.000 rpm at 4°C for a preclearing step. 100 µg for the input were saved and then cell extracts were immunoprecipitated by a GFP-Trap (Chromotek, Germany). As a negative control for pull-downs assays the eGFP plasmid (Clontech Laboratories, USA) was transfected.

For immunoprecipitation assays in CRISPR-Cas9 edited JK-G-P cells, 80 million cells were collected and lysed in ice-cold lysis buffer for 30 minutes and centrifuged for 10 minutes at 13.000 rpm at 4°C for a preclearing step. 100 µg for the input were saved. Then, steps of the GFP-Trap protocol were also followed as previously described (Castro-Sanchez et al., 2020). For Western Blot analysis of all the inputs and immunoprecipitates, solubilised cell extracts were mixed with reducing Laemmli SDS Sample Buffer 6x (Bio-Rad, USA) and boiled at 95°C for 5 minutes. Then proteins were separated by SDS-PAGE using 10% gels for resolving proteins over 50 kDa, and 13% gels for proteins under 30 kDa. Proteins were transferred to an Immobilon-FL membrane (Sigma Aldrich, USA) using a humid transfer device and 350 mA, 3 hours. Membranes were blocked for 1 hour using BSA blocking buffer (for chemiluminescence) or Li-Cor blocking buffer (for fluorescence, from Li-Cor Biosciences, USA), and incubated o/n at 4°C with the required primary antibodies. HRP-conjugated secondary goat anti-mouse or anti-rabbit (Sigma Aldrich, USA) were incubated for 1 hour at room temperature and the Pierce ECL Plus Substrate (Sigma Aldrich, USA) was used for chemiluminescence. For fluorescent Western blot IRDye 680 goat anti-rabbit and IRDye 800 goat anti-mouse (Li-Cor Biosciences, USA) were also incubated for 1 hour at RT. Both fluorescence and chemiluminescence signals were quantified using an Odyssey Infrared Imager (Li-Cor Biosciences, USA). Further, densitometry analysis was performed with the Image Studio Lite Software (version 5.2). When necessary, Blots were striped in 50 mL containing 2% SDS; 12.5% Tris-HCl pH 6.8 and 0.7%β-mercaptoethanol for 30 min at 50 ◦C.

### T cell activation and Western blot

Raji cells were pulsed with 0.5 μg/ml SEE (1 hour, 37°C, complete medium), washed and mixed with JK cells (1:10). Cells were centrifuged at 800 rpm at 37°C to promote conjugates formation. 1x10^6^ cells were lysed in ice-cold lysis buffer (5 mM Tris-HCl pH 7.5 containing 1% NP40, 0.2% Triton X-100, 150 mM NaCl, 2 mM EDTA, 1.5 mM MgCl2), and phosphatase and protease inhibitors (Sigma Aldrich, USA) followed by a preclearance step by centrifugation at 14,000 rpm (4°C, 10 min) to remove debris and nuclei. Samples were processed for SDS-PAGE with 8%-10% gels, transferred to nitrocellulose membranes using a humid transfer device at 350 mA for 3 hours, and incubated with appropriate primary and peroxidase-labelled secondary antibodies. Chemiluminescence signal was detected using the Amersham 880 detection system (GE Healthcare, Spain). When necessary, Blots were striped in 50 mL containing 2% SDS; 12.5% Tris-HCl pH 6.8 and 0.7%β-mercaptoethanol for 30 min at 50 ◦C.

### CRISPR/Cas9 genome editing

The trans-activating CRISPR RNA (tracrRNA) was mixed 1:1 with the following CRISPR RNA (crRNA): CAAGTACTGGAGCTCTGGTGG and TGAACAGCAATACAACAACCG for PRL-1 gene, ACTACACACTCACTAGAACGG and AGTTCACAGAGGTAAGATTGG for PRL-2, and GGGAAGCGATGATAACTGCGG and TCAAGCAGAGCCGGCCATACG for WDR1. Mixes were heated at 80 °C for 10 minutes and let to cool down at room temperature (RT) to facilitate the forming of the single guide RNA (sgRNA). 3.3 μM of sgRNA were incubated with Alt-R HiFi Cas9 Nuclease V3 (IDT DNA (USA) at 0.27 mg/mL and Alt-R HDR electroporation enhancer at 5 μM and then transfected into cells by using the Neon Electroporation System (Thermofisher, USA) for JK cells and the ECM 830 Square Wave Electroporation System (BTX, USA) for primary cells (in both cases the settings were 1600 V, 10 milliseconds and 3 pulses). Control cells were transfected with the nuclease duplex buffer from IDT DNA (USA) instead of crRNA.

For editing JK cells, 0.2 million cells were transfected and cultured on antibiotics-free JK medium for 24 hours. After that time, penicillin (100 U/mL) and streptomycin (100 μg/mL) were added, and cells were cultured until reaching the desired number of cells. PRL-1, PRL-2 and WDR1 edited clones were selected by limiting dilutions, until isolated clones were obtained. Due to the existence of a widely-described antibody specific for PRL-1 and PRL-2, to confirm the knockout of *PTP4A1* and *PTP4A2 genes*, protein expression was measure by Western blot. The expression of WDR1 was also evaluated by Western blot. However, *WDR1* gene was also amplified by PCR in control and WDR1 edited cells by using the following primers: forward GTAAAACGACGGCCAGT and reverse GGTCATAGCTGTTTCCTG. An expected PCR product of 261base pairs was obtained in control samples and some PCR products in control and edited samples were sequenced. The amplified DNA from the different obtained colonies were sequenced at the Genomics facility of the Complutense University of Madrid (Spain). To do so, PCR products were clone into the pGEM-T Easy Vector (Promega, Spain), transformated in DH5α competent bacteria (Thermofisher, USA), which were seeded in LB/Agar plates with 100 μg/ml ampicillin, 0.5 mM IPTG and 20 μg/ml X-Gal (all from Sigma Aldrich, USA). Plasmid was extracted by using the commercial kit from Qiagen (Germany), following manufacturers protocols.

For editing CD4 T lymphoblasts, cells were stimulated as stated before and 3 million cells were nucleofected at day 10 after isolation and cultured on in RPMI 1640 supplemented with FBS (10 %), penicillin (100 U/ml), streptomycin (100 μg/ml) and L-Glutamine (2 mM) for 72 hours.

### Immunofluorescence and confocal microscopy

For cell-cell conjugation assays, Raji B cells were labelled in serum-free medium with 10 μM 7-amino-4-chloromethylcoumarin (CMAC, Thermofisher, USA) for 1 hour at 37°C. Cells were then loaded with Staphylococcus Enterotoxin Superantigen-E (SEE, at 0.5 μg/ml, from Toxin technologies, USA) by incubation for 1 hour at 37°C, followed by two washes with complete media. Raji cells were mixed at a 1:1 ratio with JK cells and cells were spun down together to promote conjugate formation and incubated for up to 15 minutes on poly-L-lysine-coated coverslips (Sigma Aldrich, USA).

Resting JK cells or JK/Raji conjugates on coverslips were fixed with 4% paraformaldehyde (PFA, Sigma-Aldrich, USA) in TBS (NaCl 150 mM and tris-HCl pH 7.5 20 mM) during 5 minutes at room temperature (RT) in obscurity. Cells were then permeabilized with TBS-0.1% Triton X-100 (Sigma-Aldrich, USA) during 5 minutes at RT followed by blocking with 10 μg/mL human γ-globulin (Sigma-Aldrich, USA) in TBS-0.5% blocking reagent (Roche, Switzerland) for 20 minutes at RT. Samples were stained with the indicated antibodies for 1 hour at RT, washed with TBS and incubated with secondary antibodies and phalloidin-647 (Thermofisher, USA) at RT for 30 minutes in obscurity. Samples were then washed twice with TBS and once with distilled water before being mounted with Mowiol (Sigma Aldrich, USA) medium and stored at 4°C until imaging.

Confocal sections of fixed samples were acquired using a CLSM Leica TCS SP8 confocal scanning laser microscopy with a 63X/1.35 NA oil immersion objective. CMAC, Alexa 488, alexa 594 and phalloidin-647 were excited with 405, 488, 594 and 633 nm laser lines, respectively. Image acquisition was automatically optimized with the Leica software to get an image resolution of 58 nm/pixel. For 3D reconstructions, z-stacks were acquired every 0.3 μm.

### Preparation of activating lipid bilayers and TIRF microscopy

For liposomes preparation, 5 mg of DOPC and NTA lipids (Avanti Biosciences, USA) were transferred to tubes with 3 mL of chloroform (Sigma Aldrich, USA) dried under a stream of nitrogen gas. Once tubes were dry, lipids were lyophilized in a lyophilizer for 2 hours. Then, lipids were resuspended in 2 mL tris buffered saline (25mM Tris pH 8, 150mM NaCl) and kept under argon gas to prevent lipid oxidation. Multilamellar vesicles were prepared by tapping and then gentle vortexing to generate a uniform turbid suspension. Then, the multilamellar vesicles were passed through a membrane filter with an avestin extruder (0.75” diameter, 100nm pore diameter, 50X polycarbonate membrane) until the solution was clear, indicating the formation of large unilamellar vesicles. This stock was diluted 1:10 in tris buffered saline and were ready to be used.

UCHT1 anti-CD3ε Fab, recombinant ICAM-1-12His and CD58 proteins were attached to the bilayer. When necessary, UCHT1 anti-CD3ε Fab protein was labelled with Alexa 488 or 568-C5 maelamide and ICAM-1 with Alexa Fluor 405–NHS ester (all fluorophores from Thermo Scientific, USA) at ∼ 1 fluorophore per molecule ratio. For protein labelling, fluorophores (at ∼ 1 fluorophore per molecule ratio), proteins (50 μM) and TCEP (5 mM) reducing agent (Sigma Aldrich, USA) were mixed 1 hour at RT and then left overnight (O/N) at 4°C. Labelled proteins were then collected by using Zeba™Spin Desalting Columns (Thermofisher, USA).

To prepare activating lipid bilayers (ALBs), 50 μL of 12,5% NTA in DOPC large unilamellar vesicles were diluted in PBS-0.2% BSA (BSA from Sigma Aldrich) were added to ultraclean glass coverslips (Schott, USA) that were fitted with 6 channel sticky-slides (Ibidi, Germany) for 20 minutes to form supported lipid bilayers. Then, each channel was washed three times with 200 μL PBS-0.2% BSA and blocked with PBS-5% BSA for 30 minutes. Then, 100-300 molecules per μm^2^ of ICAM-1, UCHT-1 and CD58 were added for 30 minutes to perform the coating of the bilayers (Demetriou et al., 2020) After another three washes, 200.000 T cells (either JK or primary) were added to each channel and incubated at 37°C for 15 minutes. Cells were then fixed with 100 μL of PBS-4% PFA at RT for 10 minutes, washed, and permeabilized with PBS-0.1 % Triton X100. The channels were afterwards washed three times and blocked with PBS-5% BSA for 1 hour. Samples were then incubated with the desired primary antibodies for 1 hour at RT. Each channel was then washed three times with 200 μL PBS-0.2% BSA, followed by an incubation with secondary antibodies and phalloidin for another 30 minutes. The channels were washed again three times and samples were ready to be imaged.

Imaging of fixed cells over ALBs was performed with an Olympus IX83 inverted microscope (Keymed, UK) coupled to a 150X/1.45 NA oil immersion objective and an EMCCD camera (Evolve Delta, AZ) equipped with a 60× ApoN NA 1.49 objective (Olympus, JP). Alexa 405, Alexa 488, Alexa 568 and phalloidin-Alexa 647 were excited with 405, 490, 560 and 640 nm laser lines, respectively. The excitation angle was adjusted to ensure <200 nm penetration depth with respect to the basal plane. The evanescent field to image was manually selected by changing the focus into the ALBs molecules or the phalloidin staining.

### Microscopy Image analysis

Images were analysed using free software Fiji (NIH, USA). Accumulation at the IS was calculated with the Fiji plugin “Synapse measure”, as previously described (Calabia-Linares et al., 2011). Colocalisation was measured by using the built-in tool “Colocalization threshold” by manually selecting regions of interest (ROI) at the entire IS zone along z-stacks guided by Phalloidin staining. For 3D reconstructions, the Fiji tool “3D project” was used over a manually selected ROI at the IS. Actin clearance values were calculated as the ratio between the actin-free central area and the area of the phalloidin staining in the 3D reconstruction. Areas of accumulation of phalloidin and molecules over ALBs were calculated by manually doing an irregular polygon in the different channels.

### Flow cytometry

For analysis by flow cytometry of JK-Raji conjugates, Raji cells were loaded with SEE at different concentrations (by making serial dilutions 1/5 from 0.1 μg/mL to 6.4×10^-6^ μg/mL of the SEE in culture medium), incubated for 1 hour at 37 °C and washed with complete media. After this, T cells were added at a 1:1 ratio, spun together, and left overnight (o/n). Then, cell conjugates were stained for CD69 and analysed by flow cytometry. To gate out Raji cells, cell mixtures were labelled with CD19, a specific marker of B cells.

For extracellular staining, primary antibodies were incubated for 30 minutes on ice and, when needed, cells were again incubated with secondary antibodies for another 30 minutes on ice. For intracellular staining, cells were fixed for 10 minutes with PFA 5% and permeabilised for 5 minutes with 0.1% saponin in PBS (Sigma Aldrich, USA) and then blocked with PBS-FBS 2% before the staining. Next, cells were incubated on ice for 30 minutes with primary antibodies, washed and incubated for another 30 minutes with secondary antibodies and/or phalloidin. Cells were resuspended in PBS-2mM EDTA for data acquisition. A BD FACS Calibur cytometer (BD Biosciences, USA) or BD SORP LSR Fortessa (BD Biosciences, USA) were used for data acquisition. FACS data was analysed by the Flowjo software (Becton Dickinson, USA).

### ELISA

Supernatants of activated JK cells (Same cells used for activation assessment by CD69 staining) were diluted 1:50 in PBS-10% FBS and IL-2 concentration was determined by ELISA assays using the OptEIA Human IL-2 ELISA commercial kit (BD Biosciences, USA) and revealed with TMB substrate (Thermofisher, USA). Results were quantified in an ELx800 absorbance microplate reader. Absorbance values were interpolated in a standard curve using Excel software.

### Statistical analysis

All statistical analysis was implemented with PRISM8 (GraphPad software, USA). When comparing two samples, student t-test was performed. One-sample T test were used to compare normalized values to one. Comparison between three or more samples was performed with one-way ANOVA. Differences between samples along stimulation times were assess by two-way ANOVA. Specific details of each analysis are detailed in the figure legends.

## Acknowledgements

This study was supported by grants Y2018/BIO-5207_SINERGY_CAM (Madrid Regional Government) and PID2020-115444GB-I00 (MCIN/AEI/10.13039/501100011033, in part granted with ERDF A way of making Europe to PR-N. OA-S and SA-G are supported by PhD fellowships of the UCM. AG-M is supported by an Investigo Grant to NBM-C by SEPE (Fondos de Resiliencia), Gobierno de España. RR-M and PC-S are supported by SAF2012-33218 to PR-N. CCP is supported by PID2020-115444GB-I00 to PR-N and by a PhD fellowship of the UCM. MLD and SV are supported by Kennedy Trust for Rheumatology Research (202117) and European Commission ERC Horizon 2020 (SyG 951329).

