## Supplemental data for "Phosphatases of regenerating liver balance F-actin rearrangements during T cell activation through WD repeat containing protein 1"

#### Supplementary Figures and tables

**Supplementary Figure 1. SILAC proteomic approach summary. A.** Schematic of the activation approach used for the SILAC proteomics. **B**. Cofilin module proteins detected by proteomics in GFP-PRL-1 or GFP-PRL-1_D72A pull downs. The last two columns of the table indicate ratio scores of GFP PRL-1/GFP and GFP-PRL-1_D72A/GFP for the indicated proteins.

**Supplementary Figure 2. Effect of Lat-A treatment on F-actin in 293Hek cells.** Histogram of phalloidin staining obtained by flow cytometry in the indicated samples. Different samples and the mean fluorescence intensity (between brackets) are indicated.

**Supplementary Figure 3. CRISPR-Cas9 genome editing of *PTP4A1* and *PTP4A2* genes in Jurkat and primary T cells**. **A**. Schematic of the organization of *PTP4A1* and *PTP4A2* genes. The position and sequence of the crRNA guides used for genome editing are indicated. **B**. Expression of endogenous PRL-1/2 assessed by Western blot. Upper panel show expression of PRL-1/2 in control cells, polyclonal edited populations and pools of 14 *PTP4A1* and 13 *PTP4A2* selected edited clones with undetectable expression of PRL-1 or PRL-2. Lower panels show expression of PRL-1/2 in control cells and representative *PTP4A1* and *PTP4A2* edited clones. Molecular weights and the detected proteins are indicated. **C**. Graphs showing the ratio of densitometry values of PRL-1 and PRL-2 to GAPDH normalised to the control (C). The mean ± SD of the analysis of the 14 clones of *PTP4A1* edited and the 13 clones of *PTP4A2* edited is represented. **D**. Expression of endogenous PRL-1/2 assessed by Western blot of primary non-nucleofected control and edited CD4 T cells. Molecular weights and detected proteins are indicated. Right bar graphs show the ratio of the densitometry values of PRL-1 and PRL-2 and β-actin normalised to the control. Coloured symbols indicate the experiments done. Samples were compared to the control by an one-sample t-test. *$p<0.05$.

**Supplementary Figure 4. CRISPR/Cas9-based genome editing of *WDR1* gene in Jurkat and primary T cells**. **A.** Schematic of the organization of *WDR1* gene. The position and sequence of the crRNA guides used for genome editing are indicated. Lower left panel: Western blot for detection of WDR1 in control (Ctrol), polyclonal edited cells (Poly) and five representative numbered clones obtained by sorting the polyclonal population. Right graph: ratio of densitometry values of WDR1 and GAPDH normalised to the control. Colours indicate different western blot experiments and individual clones in western blot (red coloured symbols represent the data in the WB presented in the figure). **B**. Left panel: PCR to clone the DNA mutated in edited cells. Sizes in base pairs (bp) are indicated. Ladder: pair of bases ladder. Non-t: Jurkat non-transduced. Ctrol: Jurkat control. *W* ed; Jurkat *WDR1* edited. *W* C5: Jurkat *WDR1* edited clon 5. Right panels: Chromatograms of DNA sequences showing 2 examples of mutations encoding truncated WDR1 proteins and 1 example showing a mutated protein with a modified sequence. **C.** Histograms of flow cytometry of CD4 T cells edited to knockout the *WDR1* encoding gene. The isotype control and samples stained are indicated. Middle bar graphs indicate the fluorescence levels normalised to the control. Right bar graphs indicate the F-actin levels normalised to the control as detected by Phalloidin staining and flow cytometry. Symbols in graphs indicate the different donors analysed. The mean ± standard deviation is indicated. Samples were compared to the control by an one-sample t-test. *p<0.05.

**Supplementary Figure 5. Phenotype of *PTP4A1*, *PTP4A2* or *WDR1* edited JK cells**. Graphs indicate the geometric mean of fluorescence normalised to the control obtained by staining for CD3, CD4, CD28 and CD11a and flow cytometry. Samples and molecules detected are indicated. Average value and standard deviation from n=4 independent experiments (labelled by coloured symbols) are shown. Samples were compared to control JK cells with a one sample t-test. *p<0.05, ** p<0.01 and *** p<0.001. P values marginally significant are indicated.

**Supplementary Table 1: Proteomics data.** Scores of the comparison GFP-PRL-1_D72A/GFP (arranging in descending order) and GFP-PRL-1/GFP are shown. Data of WDR1, Coronin 1A and cofilin are shown bold typed.

### SUPPLEMENTARY FIGURE 1

A

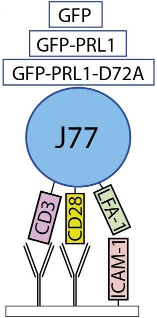


B

| Protein | Gene | Peptides | Unique peptides | Sequence coverage (%) | Molecular weight (kDa) | Intensity | Sequence lenght | Ratio GFP -PRL1/GFP | Ratio GFP-PRL1 -D72A/GFP |
|---|---|---|---|---|---|---|---|---|---|
| WDR1 | *WDR1* | 10 | 10 | 24,1 | 66,2 | 27,54 | 606 | 3,14 | 10,87 |
| PRL-1 | *PTP4A1* | 10 | 7 | 67,1 | 19,8 | 30,66 | 173 | 68,44 | 10,32 |
| Coronin 1A | *CORO1A* | 9 | 9 | 21 | 51 | 29,02 | 461 | 1,27 | 5,93 |
| Cofilin-1 | *CFL1* | 15 | 14 | 78,9 | 18,5 | 30,65 | 166 | 0,86 | 4,13 |

### SUPPLEMENTARY FIGURE 2

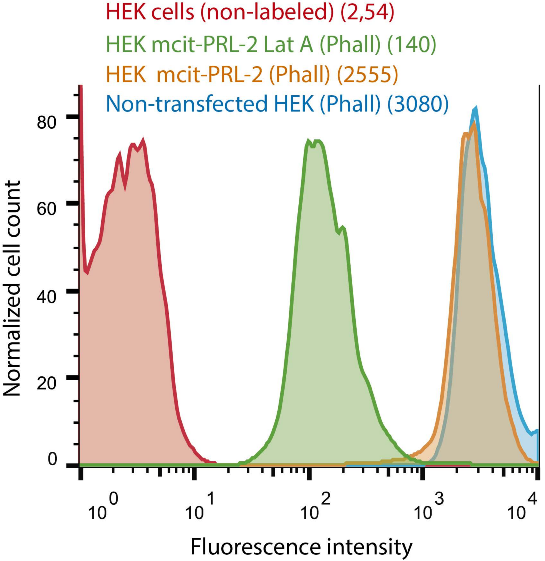

### SUPPLEMENTARY FIGURE 3

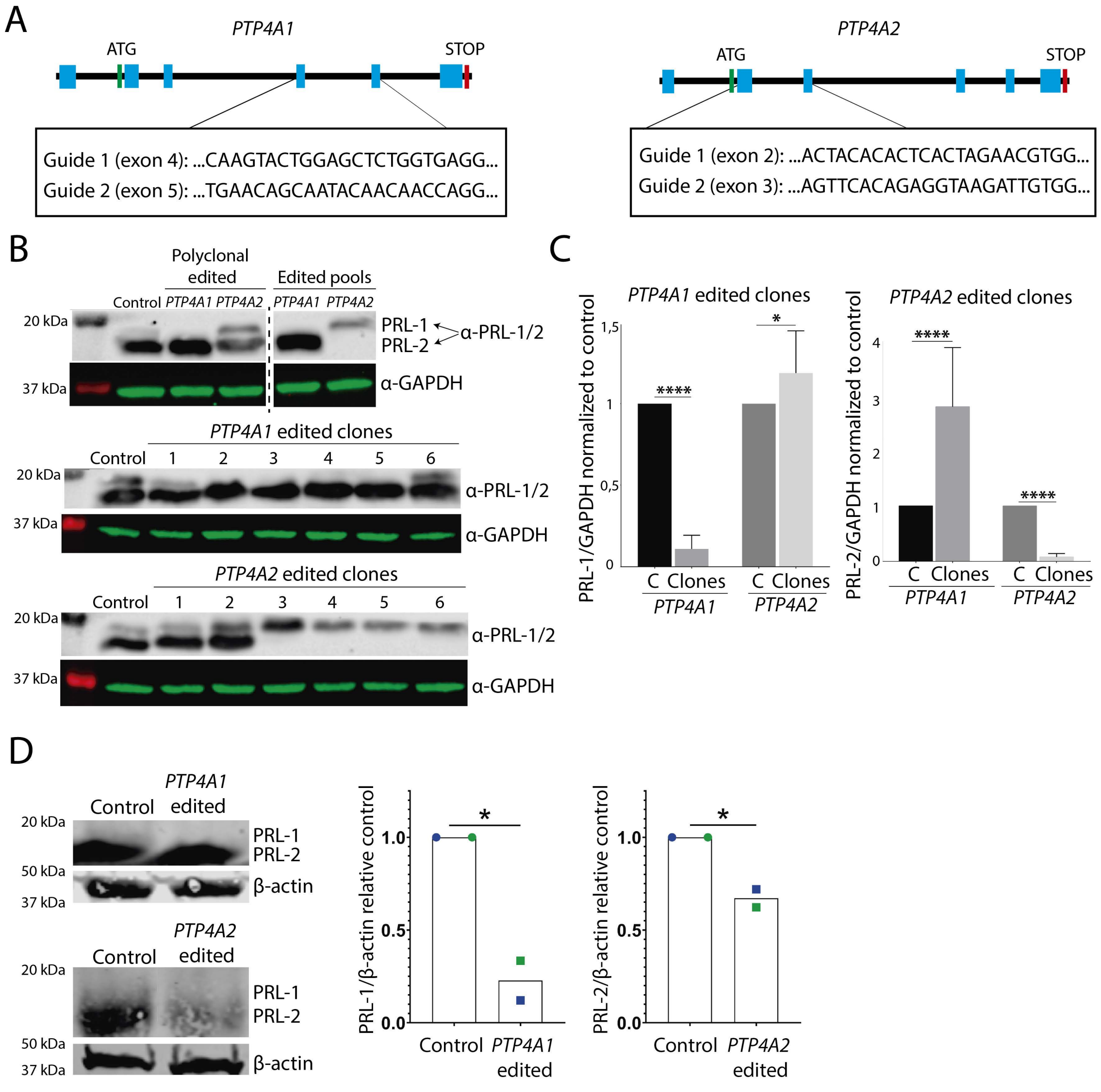

### SUPPLEMENTARY FIGURE 4

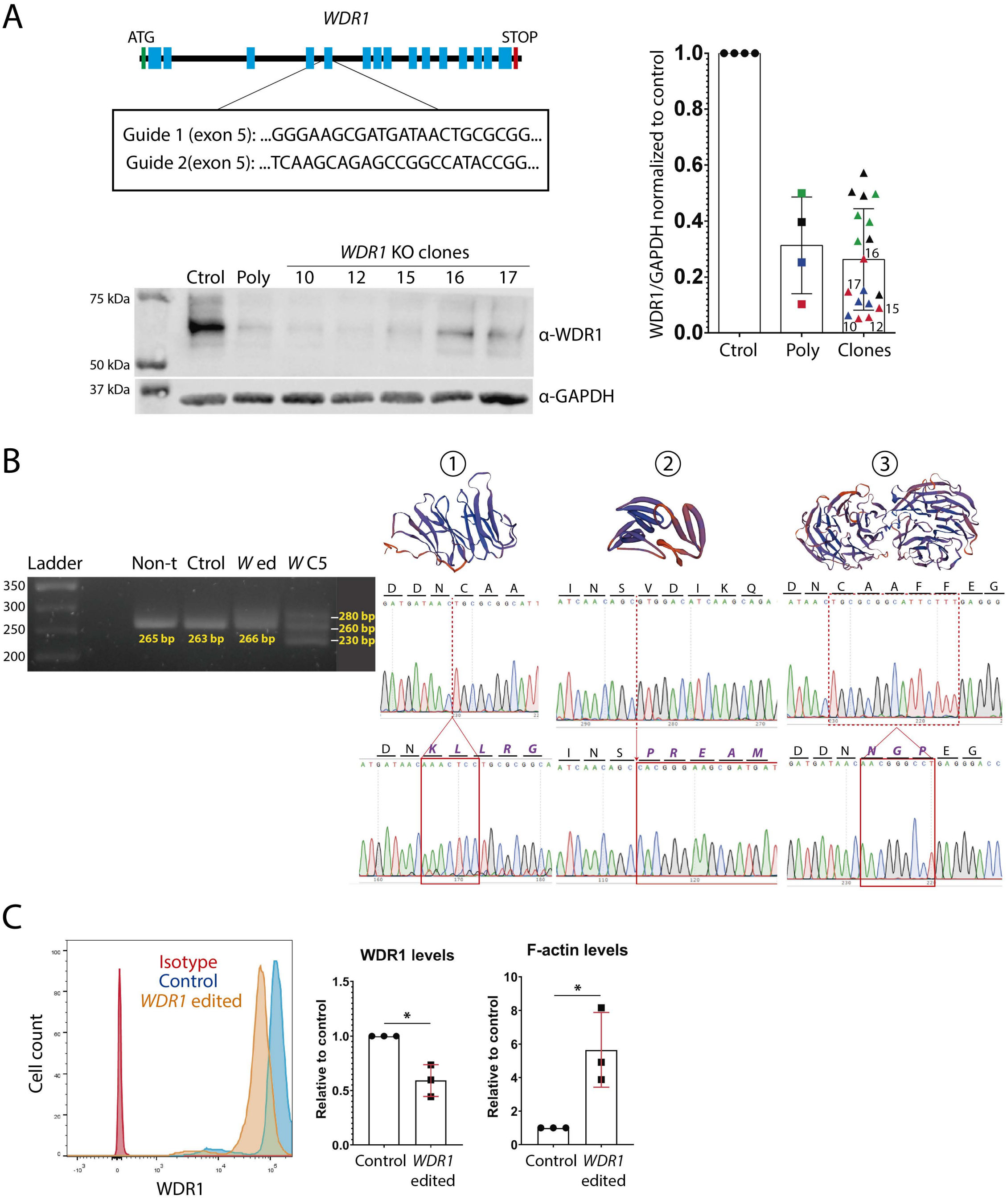

### SUPPLEMENTARY FIGURE 5

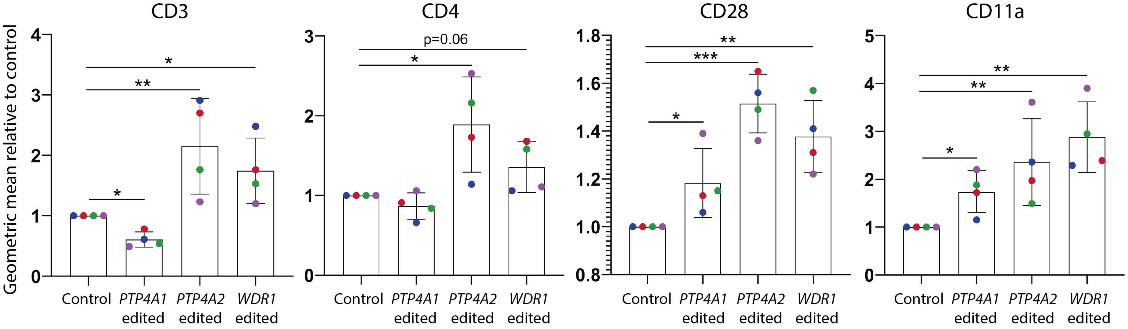

| Protein IDs | Protein names | Gene names | Peptides | Unique peptides | Sequence coverage [%] | Mol. weight [kDa] | Intensity | Sequence length | Ratio L/M D72A/GFP | Ratio H/M WT/GFP |
|---|---|---|---|---|---|---|---|---|---|---|
| P05109 | Protein S100-A | S100A8 | 6 | 6 | 47,3 | 10,8 | 27,84 | 93 | 11,51 | NaN |
| **O75083;D6** | **WD repeat** | **WDR1** | **10** | **10** | **24,1** | **66,2** | **27,54** | **606** | **10,87** | **3,14** |
| **Q93096;A0** | **Protein tyr** | **PTP4A1** | **10** | **7** | **67,1** | **19,8** | **30,66** | **173** | **10,32** | **68,44** |
| P06702 | Protein S100-A | S100A9 | 6 | 6 | 74,6 | 13,2 | 29,05 | 114 | 7,96 | 0,24 |
| P69905;G3V1 | Hemoglobin s | HBA1;HBA2 | 4 | 1 | 34,5 | 15,3 | 23,84 | 142 | 6,95 | NaN |
| P00813;F5GW | Adenosine de | ADA | 20 | 20 | 52,9 | 40,8 | 30,69 | 363 | 6,42 | 0,51 |
| **P31146;H3** | **Coronin-1A** | **CORO1A** | **9** | **9** | **21** | **51,0** | **29,02** | **461** | **5,93** | **1,27** |
| P00338;P0033 | L-lactate dehy | LDHA | 15 | 13 | 41,3 | 36,7 | 28,92 | 332 | 5,93 | 0,95 |
| Q9P258 | Protein RCC2 | RCC2 | 6 | 6 | 16,7 | 56,1 | 28,16 | 522 | 5,87 | 0,74 |
| P07900;P079C | Heat shock pr | HSP90AA1 | 41 | 29 | 54,5 | 84,7 | 30,81 | 732 | 5,63 | 1,12 |
| Q9NSD9;Q9N | Phenylalanine | FARSB | 13 | 13 | 21,4 | 66,1 | 27,27 | 589 | 5,15 | 0,91 |
| P16403 | Histone H1.2 | HIST1H1C | 8 | 3 | 32,9 | 21,4 | 26,39 | 213 | 5,05 | 1,28 |
| P63261;I3L3I0 | Actin, cytopla | ACTG1 | 23 | 1 | 64,8 | 41,8 | 29,27 | 375 | 4,87 | 0,55 |
| I3L139;Q9HA6 | Ketosamine-3- | FN3KRP | 5 | 5 | 37,2 | 15,6 | 26,73 | 145 | 4,86 | 0,66 |
| P07195;A8MV | L-lactate dehy | LDHB | 15 | 13 | 43,4 | 36,6 | 30,94 | 334 | 4,84 | 0,32 |
| Q06830;A0A0 | Peroxiredoxin- | PRDX1 | 18 | 17 | 75,9 | 22,1 | 31,60 | 199 | 4,70 | 0,65 |
| **P23528;G3** | **Cofilin-1** | **CFL1** | **15** | **14** | **78,9** | **18,5** | **30,65** | **166** | **4,13** | **0,86** |
| U3KQK0;Q998 | Histone H2B;H | HIST1H2BN;HI | 10 | 10 | 44 | 18,8 | 30,85 | 166 | 4,10 | 0,67 |
| K7EK07;Q6NX | Histone H3;Hi | H3F3B;H3F3C; | 5 | 5 | 25 | 14,9 | 28,93 | 132 | 3,92 | 0,75 |
| P07355;H0YK | Annexin A2;An | ANXA2;ANXA2 | 11 | 11 | 42,2 | 38,6 | 28,31 | 339 | 3,90 | 0,86 |
| Q16666-3;Q1 | Gamma-interf | IFI16 | 4 | 4 | 5,6 | 75,9 | 25,41 | 673 | 3,82 | 1,21 |
| P68133;P6803 | Actin, alpha sk | ACTA1;ACTC1; | 14 | 1 | 29,7 | 42,1 | 29,36 | 377 | 3,79 | 0,93 |
| Q93077;Q998 | Histone H2A t | HIST1H2AC;HI | 6 | 2 | 43,1 | 14,1 | 27,48 | 130 | 3,51 | 0,53 |
| Q15008;Q150 | 26S proteason | PSMD6 | 3 | 3 | 7,2 | 45,5 | 24,85 | 389 | 3,28 | 0,89 |
| P16401 | Histone H1.5 | HIST1H1B | 4 | 3 | 21,7 | 22,6 | 26,19 | 226 | 3,28 | 0,68 |
| P26641;P2664 | Elongation fac | EEF1G | 10 | 10 | 22,4 | 50,1 | 27,93 | 437 | 3,15 | 0,95 |
| P04406;P044C | Glyceraldehyd | GAPDH | 18 | 18 | 69,9 | 36,1 | 31,49 | 335 | 3,10 | 0,51 |
| P62805 | Histone H4 | HIST1H4A | 9 | 9 | 56,3 | 11,4 | 28,89 | 103 | 3,05 | 0,66 |
| Q08554-2;Q0 | Desmocollin-1 | DSC1 | 4 | 4 | 7,1 | 93,8 | 26,34 | 840 | 2,99 | 0,42 |
| P10599;P1059 | Thioredoxin | TXN | 5 | 5 | 40 | 11,7 | 28,34 | 105 | 2,91 | 0,74 |
| P60709;G5E9 | Actin, cytopla | ACTB | 23 | 1 | 64,8 | 41,7 | 28,79 | 375 | 2,77 | 1,03 |
| P31151;Q86S | Protein S100-A | S100A7;S100A | 6 | 6 | 67,3 | 11,5 | 27,26 | 101 | 2,76 | 0,31 |
| Q99832-3;Q9 | T-complex pro | CCT7 | 11 | 11 | 27,1 | 54,8 | 28,57 | 499 | 2,76 | 1,01 |
| P63151;P6315 | Serine/threon | PPP2R2A | 5 | 5 | 15,2 | 51,7 | 27,71 | 447 | 2,71 | 1,34 |
| P17987;E7ERF | T-complex pro | TCP1 | 22 | 22 | 45,9 | 60,3 | 30,00 | 556 | 2,60 | 1,09 |
| P52907;C9JUC | F-actin-cappin | CAPZA1 | 6 | 6 | 31,8 | 32,9 | 26,75 | 286 | 2,59 | 1,34 |
| C9J712;P3508 | Profilin-2;Prof | PFN2 | 4 | 4 | 36,3 | 9,8 | 27,04 | 91 | 2,57 | NaN |
| P40227;P4022 | T-complex pro | CCT6A | 16 | 13 | 32,4 | 58,0 | 29,13 | 531 | 2,46 | 1,23 |
| P49368;P4936 | T-complex pro | CCT3 | 17 | 17 | 42 | 60,5 | 29,12 | 545 | 2,45 | 0,90 |
| Q06323-3;Q0 | Proteasome a | PSME1 | 3 | 3 | 15,5 | 26,9 | 25,56 | 233 | 2,30 | 1,42 |
| E5RIT4;H0YBR | Eukaryotic tra | EIF3E | 2 | 2 | 26,7 | 8,9 | 26,49 | 75 | 2,23 | 0,95 |
| A0A087WW6 | 26S proteason | PSMD1 | 5 | 5 | 5,9 | 105,9 | 25,32 | 953 | 2,21 | 0,92 |
| P13639 | Elongation fac | EEF2 | 10 | 10 | 10,8 | 95,3 | 27,39 | 858 | 2,19 | 1,11 |
| P08238;Q58F | Heat shock pr | HSP90AB1 | 32 | 16 | 47 | 83,3 | 29,39 | 724 | 2,11 | 0,57 |
| P68871;F8W6 | Hemoglobin s | HBB;HBD | 6 | 5 | 48,3 | 16,0 | 26,32 | 147 | 2,10 | 0,81 |
| E9PBS1;P2223 | Multifunction | PAICS | 8 | 8 | 17,4 | 45,7 | 28,28 | 413 | 2,07 | 0,77 |
| P06753-2;Q5H | Tropomyosin a | TPM3;DKFZp6 | 9 | 9 | 46,4 | 29,0 | 26,90 | 248 | 2,05 | 0,74 |
| E9PL71;E9PQ4 | Elongation fac | EEF1D | 5 | 5 | 29,4 | 20,8 | 26,38 | 187 | 1,99 | 0,84 |
| P50990;P5099 | T-complex pro | CCT8 | 21 | 21 | 39,8 | 59,6 | 29,13 | 548 | 1,97 | 1,66 |
| E7ENZ3;P4864 | T-complex pro | CCT5 | 14 | 14 | 32,1 | 53,8 | 29,22 | 486 | 1,96 | 1,43 |
| P50991;P5099 | T-complex pro | CCT4 | 18 | 18 | 38,6 | 57,9 | 29,47 | 539 | 1,93 | 0,97 |
| P62857 | 40S ribosomal | RPS28 | 4 | 4 | 58 | 7,8 | 28,06 | 69 | 1,91 | 0,46 |
| Q14152-2;Q1 | Eukaryotic tra | EIF3A | 6 | 6 | 7,6 | 162,6 | 25,82 | 1348 | 1,83 | 0,95 |
| P40939;H0YF | Trifunctional e | HADHA | 7 | 7 | 12,2 | 83,0 | 26,29 | 763 | 1,82 | 1,26 |
| E7EQB2;E7ER | Lactotransferr | LTF | 4 | 4 | 6,2 | 76,6 | 23,98 | 696 | 1,76 | 0,79 |
| Q13263-2;Q1 | Transcription i | TRIM28 | 7 | 7 | 13,3 | 79,5 | 27,59 | 753 | 1,73 | 1,04 |
| K7EMD0;K7EP | Proteasome a | PSME3 | 2 | 2 | 26 | 10,9 | 24,14 | 96 | 1,72 | 1,10 |
| A0A087WZK9; | Eukaryotic tra | EIF3H | 2 | 2 | 8 | 39,6 | 26,45 | 349 | 1,71 | 1,17 |
| P35579;P3557 | Myosin-9 | MYH9 | 26 | 24 | 19,5 | 226,5 | 28,90 | 1960 | 1,70 | 0,94 |
| Q15365 | Poly(rC)-bindi | PCBP1 | 8 | 4 | 27,8 | 37,5 | 26,11 | 356 | 1,70 | 1,26 |
| B4DY09;Q129 | Interleukin en | ILF2 | 3 | 3 | 12,8 | 38,9 | 25,96 | 352 | 1,60 | 0,75 |
| P01040;C9J0E | Cystatin-A;Cys | CSTA | 6 | 6 | 75,5 | 11,0 | 25,73 | 98 | 1,60 | 1,61 |
| C9JXK0;Q1473 | Lamin-B recep | LBR | 2 | 2 | 9,4 | 24,3 | 26,68 | 213 | 1,58 | 1,37 |
| Q8NC51-4;Q8 | Plasminogen a | SERBP1 | 6 | 6 | 16,3 | 42,4 | 26,66 | 387 | 1,58 | 0,60 |
| P12273 | Prolactin-indu | PIP | 3 | 3 | 32,9 | 16,6 | 24,34 | 146 | 1,57 | NaN |
| P25398 | 40S ribosomal | RPS12 | 7 | 7 | 68,9 | 14,5 | 29,10 | 132 | 1,56 | 0,67 |
| B1AHC9;P129 | X-ray repair cr | XRCC6 | 8 | 8 | 15,7 | 64,3 | 26,37 | 559 | 1,53 | 1,07 |
| P35998;C9JLS | 26S protease r | PSMC2 | 6 | 6 | 16,4 | 48,6 | 26,20 | 433 | 1,53 | 1,25 |
| F8VXL2;F8VR | Dynein light ch | DYNLL1 | 2 | 2 | 17 | 5,4 | 25,82 | 47 | 1,50 | 1,20 |
| F8W1R7;G3V1 | Myosin light p | MYL6 | 6 | 6 | 41,4 | 16,3 | 27,68 | 145 | 1,49 | 1,29 |
| F8VQ14;F5GW | T-complex pro | CCT2 | 16 | 16 | 44,7 | 44,8 | 28,57 | 416 | 1,49 | 1,18 |
| P22626-2;P22 | Heterogeneou | HNRNPA2B1 | 13 | 13 | 45,7 | 36,0 | 28,25 | 341 | 1,48 | 0,77 |
| Q9UBN7;C9J1 | Histone deace | HDAC6 | 8 | 8 | 6,3 | 131,4 | 27,88 | 1215 | 1,47 | 0,67 |

| | | | | | | | | | |
|---|---|---|---|---|---|---|---|---|---|
| Q09028-4;Q09 | Histone-bindir | RBBP4;RBBP7 | 2 | 2 | 7,4 | 43,5 | 23,75 | 390 | 1,46 | 0,83 |
| P06748-3;P06 | Nucleophosmi | NPM1 | 9 | 9 | 37,8 | 28,4 | 29,25 | 259 | 1,45 | 0,92 |
| P21796;C9JI87 | Voltage-deper | VDAC1 | 14 | 14 | 58 | 30,8 | 28,98 | 283 | 1,45 | 1,88 |
| G5E972;P4216 | Lamina-associ | TMPO | 8 | 8 | 25,1 | 46,3 | 26,71 | 414 | 1,41 | 1,43 |
| O15145;C9JZD | Actin-related | ARPC3 | 7 | 7 | 41,6 | 20,5 | 27,37 | 178 | 1,40 | 0,59 |
| F8W6I7;P0965 | Heterogeneou | HNRNPA1;HNI | 12 | 11 | 35,5 | 33,2 | 27,66 | 307 | 1,38 | 0,81 |
| O43175;Q5SZI | D-3-phosphog | PHGDH | 13 | 13 | 29,8 | 56,7 | 32,77 | 533 | 1,38 | 1,16 |
| Q9NR30-2;Q9 | Nucleolar RNA | DDX21;DDX50 | 2 | 2 | 3,1 | 79,7 | 25,20 | 715 | 1,37 | 0,85 |
| H3BTN5;P146 | Pyruvate kina | PKM | 10 | 10 | 29,5 | 53,0 | 27,86 | 485 | 1,36 | 0,79 |
| P38159;H3BT | RNA-binding n | RBMX;RBMXL | 6 | 6 | 18,9 | 42,3 | 25,57 | 391 | 1,36 | 0,69 |
| P68363;P6836 | Tubulin alpha- | TUBA1B;TUBA | 16 | 16 | 44,1 | 50,2 | 30,94 | 451 | 1,36 | 1,12 |
| P62979 | Ubiquitin-40S | RPS27A | 5 | 2 | 46,2 | 18,0 | 23,87 | 156 | 1,34 | 0,78 |
| F8W7C6;A0A0 | 60S ribosomal | RPL10 | 4 | 4 | 25,8 | 18,6 | 26,31 | 163 | 1,31 | 0,95 |
| B3KM87;A0A0 | Matrin-3 | MATR3;DKFZp | 2 | 2 | 5,1 | 56,7 | 24,35 | 509 | 1,29 | 0,98 |
| F8VWS0;P053 | 60S acidic ribc | RPLP0;RPLP0P | 7 | 7 | 35,9 | 30,5 | 27,10 | 281 | 1,29 | 1,17 |
| Q9UMS4;H0Y | Pre-mRNA-prc | PRPF19 | 5 | 5 | 8,1 | 55,2 | 25,82 | 504 | 1,29 | 1,07 |
| Q9UQ35;Q9U | Serine/arginin | SRRM2 | 7 | 7 | 4 | 299,6 | 26,24 | 2752 | 1,29 | 0,67 |
| H0YKD8;P467 | 60S ribosomal | RPL28 | 3 | 3 | 17,6 | 19,1 | 25,54 | 170 | 1,28 | 1,02 |
| J3KTJ8;J3QRI | 60S ribosomal | RPL26;RPL26L | 4 | 4 | 42,7 | 11,5 | 25,62 | 96 | 1,28 | 0,95 |
| P43487-2;P43 | Ran-specific G | RANBP1 | 4 | 4 | 30 | 23,2 | 24,87 | 200 | 1,28 | 0,96 |
| P00558-2;P00 | Phosphoglyce | PGK1 | 11 | 11 | 38 | 41,4 | 27,65 | 389 | 1,28 | NaN |
| G3V279;P840 | Enhancer of ru | ERH | 1 | 1 | 15,5 | 8,2 | 24,67 | 71 | 1,27 | 0,31 |
| M0R181;P467 | 60S ribosomal | RPL21 | 3 | 3 | 23,8 | 14,1 | 25,71 | 122 | 1,27 | 1,02 |
| J3KR24;A0A0 | Isoleucine--tRI | IARS | 4 | 4 | 5,2 | 131,8 | 25,50 | 1152 | 1,27 | 1,02 |
| P61978-3;P61 | Heterogeneou | HNRNPK | 5 | 5 | 12,7 | 48,6 | 26,96 | 440 | 1,27 | 0,93 |
| Q9H307;Q9H3 | Pinin | PNN | 3 | 3 | 4,9 | 81,6 | 24,79 | 717 | 1,24 | 0,81 |
| Q5JP53;P0743 | Tubulin beta c | TUBB | 17 | 3 | 51,2 | 47,8 | 28,15 | 426 | 1,23 | 1,06 |
| P36578;H3BT | 60S ribosomal | RPL4 | 5 | 5 | 13,8 | 47,7 | 28,13 | 427 | 1,23 | 0,90 |
| Q8N5F7;Q5M | NF-kappa-B-a | NKAP;NKAPL | 2 | 2 | 6 | 47,1 | 24,32 | 415 | 1,23 | 1,00 |
| E9PKZ0;P6291 | 60S ribosomal | RPL8 | 5 | 5 | 20,5 | 22,4 | 25,84 | 205 | 1,23 | 0,90 |
| Q12906-5;Q1 | Interleukin en | ILF3 | 4 | 4 | 7,1 | 74,6 | 23,79 | 690 | 1,22 | 1,06 |
| Q9Y277;Q9Y2 | Voltage-deper | VDAC3 | 7 | 7 | 37,5 | 30,7 | 27,31 | 283 | 1,20 | 1,46 |
| A0A087WUK2 | Heterogeneou | HNRNPDL | 3 | 2 | 10,2 | 40,0 | 25,79 | 363 | 1,20 | 0,61 |
| P46777;R4GN | 60S ribosomal | RPL5 | 9 | 9 | 29,6 | 34,4 | 28,30 | 297 | 1,19 | 0,94 |
| H0YEN5;P158 | 40S ribosomal | RPS2 | 5 | 5 | 30,8 | 21,2 | 26,05 | 195 | 1,19 | 0,99 |
| G3V4W0;B4D | Heterogeneou | HNRNPC | 15 | 15 | 43,9 | 28,9 | 28,72 | 262 | 1,19 | 0,88 |
| A0A087X0X3; | Heterogeneou | HNRNPM | 11 | 11 | 15,3 | 77,6 | 28,07 | 730 | 1,18 | 1,10 |
| P61353;K7ER | 60S ribosomal | RPL27 | 2 | 2 | 11,8 | 15,8 | 24,70 | 136 | 1,17 | 1,11 |
| P05023-4;P05 | Sodium/potas | ATP1A1;ATP1 | 12 | 9 | 14,3 | 113,0 | 25,98 | 1023 | 1,17 | 1,26 |
| P19338;H7BY | Nucleolin | NCL | 22 | 22 | 29,7 | 76,6 | 29,61 | 710 | 1,17 | 0,82 |
| P62424;Q5T8 | 60S ribosomal | RPL7A | 12 | 12 | 38 | 30,0 | 27,61 | 266 | 1,16 | 0,89 |
| Q14204 | Cytoplasmic d | DYNC1H1 | 11 | 11 | 3,3 | 532,4 | 26,70 | 4646 | 1,16 | 0,96 |
| P45880-2;P45 | Voltage-deper | VDAC2 | 12 | 12 | 44,9 | 30,4 | 28,15 | 283 | 1,16 | 1,60 |
| G3V2C9;P597 | Guanine nucle | GNG2 | 1 | 1 | 25 | 5,9 | 23,29 | 52 | 1,16 | 1,19 |
| K7EQJ5;P6284 | 40S ribosomal | RPS15 | 3 | 3 | 22,7 | 16,6 | 27,38 | 141 | 1,16 | 1,25 |
| A0A087WXM | 60S ribosomal | RPL17;RPL17- | 6 | 6 | 42 | 19,6 | 27,77 | 169 | 1,15 | 0,89 |
| P39023;G5E9 | 60S ribosomal | RPL3 | 11 | 10 | 23,3 | 46,1 | 28,12 | 403 | 1,15 | 0,92 |
| P31930;G3V0 | Cytochrome b | UQCRC1 | 6 | 6 | 16,7 | 52,6 | 27,89 | 480 | 1,14 | 3,00 |
| M0QXM4;Q15 | Amino acid tra | SLC1A5 | 1 | 1 | 4,1 | 39,4 | 25,78 | 365 | 1,13 | 0,94 |
| Q07666-2;Q0 | KH domain-co | KHDRBS1;KHD | 2 | 2 | 4,5 | 45,9 | 25,47 | 418 | 1,13 | 1,12 |
| Q92901 | 60S ribosomal | RPL3L | 3 | 2 | 4,4 | 46,3 | 25,85 | 407 | 1,13 | 1,03 |
| P62906 | 60S ribosomal | RPL10A | 5 | 5 | 26,3 | 24,8 | 27,03 | 217 | 1,12 | 0,93 |
| H0Y449;P678 | Nuclease-sens | YBX1 | 6 | 6 | 26,5 | 41,9 | 26,08 | 374 | 1,12 | 0,79 |
| P07814;V9GY | Bifunctional gl | EPRS | 14 | 14 | 10,5 | 170,6 | 27,72 | 1512 | 1,11 | 0,83 |
| C9JYS8;Q1523 | Non-POU dom | NONO | 3 | 3 | 12,6 | 29,5 | 24,54 | 247 | 1,11 | 1,22 |
| P23246-2;P23 | Splicing factor | SFPQ | 3 | 3 | 3,4 | 72,3 | 25,11 | 669 | 1,10 | 0,81 |
| Q5VTE0;P681 | Putative elong | EEF1A1P5;EEF | 21 | 21 | 50,6 | 50,2 | 31,72 | 462 | 1,10 | 1,05 |
| C9JXB8;C9JN | 60S ribosomal | RPL24 | 3 | 3 | 24 | 14,4 | 25,89 | 121 | 1,10 | 1,03 |
| K7EMA7;H7B | 60S ribosomal | RPL23A | 4 | 4 | 47,1 | 7,9 | 26,83 | 70 | 1,09 | 1,04 |
| P18077;C9K0 | 60S ribosomal | RPL35A | 5 | 5 | 35,5 | 12,5 | 27,21 | 110 | 1,09 | 0,92 |
| P62753;A2A3 | 40S ribosomal | RPS6 | 8 | 8 | 24,5 | 28,7 | 28,11 | 249 | 1,08 | 0,83 |
| P62081;B5MC | 40S ribosomal | RPS7 | 4 | 4 | 25,3 | 22,1 | 24,97 | 194 | 1,08 | 0,99 |
| Q00839;Q008 | Heterogeneou | HNRNPU | 12 | 12 | 16,8 | 90,6 | 28,48 | 825 | 1,07 | 0,88 |
| F6S8N6;A0A0 | Protein-L-isoa | PCMT1 | 3 | 3 | 12,8 | 26,6 | 27,47 | 250 | 1,07 | 0,69 |
| Q5JR95;P6224 | 40S ribosomal | RPS8 | 8 | 8 | 39,9 | 21,9 | 27,93 | 188 | 1,06 | 0,88 |
| P53621;P5362 | Coatomer sub | COPA | 3 | 3 | 3,4 | 138,3 | 23,34 | 1224 | 1,06 | 0,68 |
| P61247;D6RA | 40S ribosomal | RPS3A | 8 | 8 | 33,7 | 29,9 | 27,98 | 264 | 1,06 | 1,01 |
| P08670;B0YJC | Vimentin | VIM | 21 | 18 | 53 | 53,7 | 29,34 | 466 | 1,06 | 1,32 |
| P38646;D6RJI | Stress-70 prot | HSPA9 | 29 | 29 | 48,5 | 73,7 | 29,96 | 679 | 1,05 | 0,99 |
| H0YD64;E9PS | Solute carrier | SLC43A3 | 1 | 1 | 9,5 | 15,7 | 24,40 | 137 | 1,05 | 1,36 |
| P26373;P2637 | 60S ribosomal | RPL13 | 6 | 6 | 28 | 24,3 | 28,75 | 211 | 1,04 | 0,91 |
| C9JW96;P352 | Prohibitin | PHB | 7 | 7 | 25,2 | 26,9 | 27,95 | 246 | 1,04 | 1,38 |
| P18124;A8MU | 60S ribosomal | RPL7 | 3 | 3 | 16,5 | 29,2 | 26,57 | 248 | 1,04 | 0,74 |
| D3YTB1;F8W7 | 60S ribosomal | RPL32 | 3 | 3 | 23,3 | 15,6 | 26,25 | 133 | 1,03 | 0,96 |
| Q9UHD1-2;Q9 | Cysteine and | CHORDC1 | 4 | 4 | 20,1 | 35,3 | 24,80 | 313 | 1,02 | 0,28 |
| Q9P0L0;Q9P0 | Vesicle-associ | VAPA | 8 | 7 | 37,8 | 27,9 | 28,13 | 249 | 1,00 | 1,28 |

| | | | | | | | | | | |
|---|---|---|---|---|---|---|---|---|---|---|
| E7EN40;D6RIL | Heterogeneou | HNRNPH1;HNI | 3 | 3 | 26,5 | 18,8 | 25,57 | 166 | 1,00 | 0,76 |
| H0Y5B4;H7BZ: | 60S ribosomal | RPL36A;RPL36 | 3 | 3 | 30,4 | 13,2 | 26,05 | 112 | 1,00 | 0,82 |
| Q9Y266;A0A0. | Nuclear migra | NUDC | 8 | 8 | 27,8 | 38,2 | 27,33 | 331 | 1,00 | 1,48 |
| P61224-2;P61 | Ras-related pr | RAP1B;RAP1A | 5 | 5 | 38,7 | 15,4 | 25,27 | 137 | 1,00 | 0,96 |
| Q86VP6;Q86V | Cullin-associat | CAND1 | 10 | 10 | 12,2 | 136,4 | 26,95 | 1230 | 1,00 | 0,53 |
| P20700;E9PBF | Lamin-B1 | LMNB1 | 21 | 20 | 43,7 | 66,4 | 28,44 | 586 | 1,00 | 1,18 |
| P25705;P257C | ATP synthase : | ATP5A1 | 22 | 22 | 46,3 | 59,8 | 30,87 | 553 | 0,99 | 1,07 |
| P51572;P5157 | B-cell receptor | BCAP31 | 7 | 7 | 26 | 28,0 | 26,48 | 246 | 0,99 | 1,33 |
| M0R0P7;M0R: | 60S ribosomal | RPL18A | 4 | 4 | 19,7 | 16,2 | 26,41 | 137 | 0,98 | 0,85 |
| A0A087WTT1; | Polyadenylate | PABPC1;PABP( | 3 | 2 | 6,9 | 58,5 | 23,79 | 522 | 0,98 | 0,93 |
| P61313;E7EQ\ | 60S ribosomal | RPL15 | 7 | 7 | 27,9 | 24,1 | 26,74 | 204 | 0,98 | 0,98 |
| P60866;P608€ | 40S ribosomal | RPS20 | 7 | 7 | 45,4 | 13,4 | 29,66 | 119 | 0,98 | 0,82 |
| C9JRZ6;Q9NX( | MICOS comple | CHCHD3 | 8 | 8 | 32,3 | 26,7 | 27,72 | 232 | 0,98 | 1,52 |
| Q07020;F8VU. | 60S ribosomal | RPL18 | 7 | 7 | 37,2 | 21,6 | 27,22 | 188 | 0,97 | 0,94 |
| P13073;Q86W | Cytochrome c | COX4I1 | 7 | 7 | 39,1 | 19,6 | 29,10 | 169 | 0,97 | 4,56 |
| Q02878;F8VZ4 | 60S ribosomal | RPL6 | 11 | 11 | 32,3 | 32,7 | 28,12 | 288 | 0,97 | 0,95 |
| P40429;Q8J01 | 60S ribosomal | RPL13A;RPL13 | 4 | 4 | 23,2 | 23,6 | 27,88 | 203 | 0,96 | 0,99 |
| I3L3H2;P3891! | Eukaryotic init | EIF4A3 | 2 | 2 | 22,4 | 13,6 | 26,13 | 125 | 0,95 | 1,03 |
| P22695;H3BR( | Cytochrome b | UQCRC2 | 8 | 8 | 22,7 | 48,4 | 28,59 | 453 | 0,94 | 3,39 |
| P46783;F6U21 | 40S ribosomal | RPS10;RPS10-I | 4 | 4 | 20,6 | 18,9 | 27,30 | 165 | 0,94 | 1,16 |
| K7ERG4;P623: | Small nuclear | SNRPD2 | 2 | 2 | 21,8 | 8,8 | 25,62 | 78 | 0,93 | 0,73 |
| P62263;H0YB2 | 40S ribosomal | RPS14 | 5 | 5 | 43,7 | 16,3 | 28,06 | 151 | 0,93 | 0,86 |
| E5RJR5;P6320 | S-phase kinase | SKP1 | 1 | 1 | 8 | 18,7 | 24,36 | 163 | 0,93 | 0,68 |
| Q5QNZ2;P245 | ATP synthase | ATP5F1 | 4 | 4 | 22,1 | 22,3 | 27,65 | 195 | 0,92 | 0,97 |
| P11142;E9PKE | Heat shock co | HSPA8 | 33 | 28 | 55,4 | 70,9 | 31,08 | 646 | 0,92 | 0,98 |
| E9PLX7;P4677 | 60S ribosomal | RPL27A | 3 | 3 | 28,3 | 12,5 | 27,33 | 113 | 0,91 | 0,83 |
| J3QR09;J3KTE | Ribosomal pro | RPL19 | 6 | 6 | 27,5 | 23,1 | 26,94 | 193 | 0,90 | 1,13 |
| Q96AG4 | Leucine-rich re | LRRC59 | 8 | 8 | 23,8 | 34,9 | 28,16 | 307 | 0,89 | 1,02 |
| P08708;H0YN! | 40S ribosomal | RPS17 | 5 | 5 | 40 | 15,6 | 26,60 | 135 | 0,89 | 1,09 |
| P61981;Q4VY: | 14-3-3 protein | YWHAG;YWH/ | 4 | 3 | 13,8 | 28,3 | 25,13 | 247 | 0,87 | 1,04 |
| H0YA96;H0Y8( | Heterogeneou | HNRNPD | 4 | 3 | 20,5 | 23,8 | 26,04 | 210 | 0,87 | 0,69 |
| P14625;Q96G! | Endoplasmin | HSP90B1 | 7 | 5 | 7,8 | 92,5 | 26,51 | 803 | 0,86 | 0,93 |
| A0A087WYT3; | Prostaglandin | PTGES3 | 2 | 2 | 18,3 | 19,2 | 24,68 | 164 | 0,86 | 0,68 |
| P62266;D6RD | 40S ribosomal | RPS23 | 3 | 3 | 21 | 15,8 | 27,86 | 143 | 0,85 | 0,81 |
| Q9UJV9;J3KNI | Probable ATP- | DDX41 | 4 | 4 | 6,3 | 69,8 | 24,10 | 622 | 0,84 | 0,98 |
| P27797;K7EJB | Calreticulin | CALR | 8 | 8 | 23 | 48,1 | 27,76 | 417 | 0,84 | 1,61 |
| B9A067;Q168! | MICOS comple | IMMT | 19 | 19 | 27,8 | 79,0 | 28,88 | 711 | 0,84 | 1,30 |
| J3KPX7;Q9962 | Prohibitin-2 | PHB2 | 6 | 6 | 21,1 | 33,2 | 28,23 | 298 | 0,84 | 1,35 |
| H0YCP8;A0A0. | Poly(U)-bindin | PUF60 | 2 | 2 | 7,7 | 28,1 | 23,63 | 273 | 0,84 | 0,64 |
| P62829;J3KTJ: | 60S ribosomal | RPL23 | 3 | 3 | 18,6 | 14,9 | 27,81 | 140 | 0,82 | 0,88 |
| I3L159;I3L1F5, | Heme oxygena | HMOX2 | 2 | 2 | 10,9 | 26,6 | 24,15 | 229 | 0,82 | 1,00 |
| P55060-3;P55 | Exportin-2 | CSE1L | 13 | 13 | 16,6 | 107,8 | 27,10 | 945 | 0,82 | 1,02 |
| P05387;H0YDI | 60S acidic ribc | RPLP2 | 3 | 3 | 53 | 11,7 | 28,07 | 115 | 0,82 | 1,30 |
| H3BMH2;H3B! | Ras-related pr | RAB11A;RAB1 | 5 | 5 | 33,5 | 17,7 | 27,29 | 155 | 0,81 | 1,18 |
| P84095 | Rho-related G | RHOG | 2 | 2 | 15,7 | 21,3 | 24,87 | 191 | 0,81 | 1,02 |
| B1AH77;B1AH | Ras-related C3 | RAC2;RAC1;R/ | 4 | 4 | 29,7 | 16,8 | 27,54 | 148 | 0,81 | 1,22 |
| P62701;C9JEH | 40S ribosomal | RPS4X | 8 | 8 | 33,5 | 29,6 | 27,46 | 263 | 0,81 | 0,84 |
| G5E9R3;Q6P4 | 60S ribosomal | RPL37A | 3 | 3 | 41,4 | 6,6 | 26,58 | 58 | 0,81 | 0,90 |
| P10809;E7ESH | 60 kDa heat sh | HSPD1 | 41 | 41 | 68,1 | 61,1 | 33,07 | 573 | 0,80 | 1,24 |
| P62851;E9PQI | 40S ribosomal | RPS25 | 5 | 5 | 33,6 | 13,7 | 27,91 | 125 | 0,80 | 0,82 |
| Q15125 | 3-beta-hydrox | EBP | 1 | 1 | 5,2 | 26,4 | 26,05 | 230 | 0,79 | 1,12 |
| C9J4V0;C9J8S: | Ras-related pr | RAB7A | 6 | 6 | 49,3 | 15,1 | 27,67 | 134 | 0,79 | 1,24 |
| P05386 | 60S acidic ribc | RPLP1 | 2 | 2 | 36,8 | 11,5 | 25,26 | 114 | 0,79 | 0,86 |
| Q96HJ9-2;A0A | Putative RNA- | LUC7L2 | 3 | 3 | 8,5 | 54,2 | 24,48 | 458 | 0,78 | 0,49 |
| A0A087WYB4; | Stomatin-like | STOML2 | 8 | 8 | 30 | 33,4 | 27,24 | 310 | 0,78 | 1,42 |
| P61006;P610C | Ras-related pr | RAB8A | 5 | 4 | 29 | 23,7 | 25,61 | 207 | 0,78 | 1,26 |
| P62269;J3JS6! | 40S ribosomal | RPS18 | 3 | 3 | 17,1 | 17,7 | 24,55 | 152 | 0,76 | 0,70 |
| P27824;P2782 | Calnexin | CANX | 13 | 13 | 24,3 | 67,6 | 27,98 | 592 | 0,76 | 1,59 |
| P49755;G3V2I | Transmembra | TMED10 | 3 | 3 | 16,4 | 25,0 | 25,79 | 219 | 0,76 | 1,19 |
| P16615-5;P16 | Sarcoplasmic/ | ATP2A2 | 8 | 8 | 10,6 | 109,7 | 26,98 | 997 | 0,75 | 1,11 |
| X6R433;A0A0/ | Protein-tyrosi | PTPRC | 3 | 3 | 2,5 | 131,1 | 24,01 | 1145 | 0,74 | 1,48 |
| Q8TA86;C9J6\ | Retinitis pigm | RP9 | 6 | 6 | 30,8 | 26,1 | 28,39 | 221 | 0,73 | 0,84 |
| P11021 | 78 kDa glucos | HSPA5 | 15 | 14 | 27,2 | 72,3 | 27,83 | 654 | 0,73 | 0,84 |
| J3KS45;J9JIE6; | Transmembra | TMCO1 | 2 | 2 | 14 | 16,6 | 25,73 | 143 | 0,73 | 1,34 |
| C9JAW5;C9JN | HIG1 domain f | HIGD1A | 1 | 1 | 21,7 | 8,8 | 23,47 | 83 | 0,73 | 1,03 |
| Q9NWB6;Q9N | Arginine and g | ARGLU1 | 11 | 11 | 29,3 | 33,2 | 29,80 | 273 | 0,73 | 0,80 |
| E9PCX7;Q134 | NAD(P) transh | NNT | 3 | 3 | 3,4 | 99,7 | 24,55 | 955 | 0,73 | 1,27 |
| P49411;H3BN | Elongation fac | TUFM | 6 | 6 | 18,1 | 49,5 | 27,09 | 452 | 0,72 | 1,17 |
| X6RFL8;P6110 | Ras-related pr | RAB14 | 2 | 2 | 8,3 | 20,4 | 25,12 | 181 | 0,72 | 0,63 |
| A0A087WVQ6 | Clathrin heavy | CLTC | 8 | 8 | 5,4 | 192,1 | 25,92 | 1679 | 0,71 | 1,04 |
| P04843;B7Z4L | Dolichyl-diphc | RPN1 | 11 | 11 | 19,4 | 68,6 | 28,73 | 607 | 0,71 | 1,07 |
| P62273;P6227 | 40S ribosomal | RPS29 | 3 | 3 | 33,9 | 6,7 | 24,83 | 56 | 0,70 | 0,74 |
| Q92575 | UBX domain-c | UBXN4 | 3 | 3 | 4,9 | 56,8 | 25,11 | 508 | 0,70 | 1,28 |
| C9JDR0;Q157: | Sterol-4-alpha | NSDHL | 3 | 3 | 12,2 | 28,1 | 26,17 | 254 | 0,69 | 1,14 |
| O75947;O759 | ATP synthase : | ATP5H | 6 | 6 | 37,9 | 18,5 | 27,65 | 161 | 0,69 | 1,06 |
| P39019;A0A0 | 40S ribosomal | RPS19 | 7 | 7 | 42,8 | 16,1 | 27,17 | 145 | 0,68 | 1,07 |

| | | | | | | | | | | |
|---|---|---|---|---|---|---|---|---|---|---|
| F5GZS6;J3KPF | 4F2 cell-surfac | SLC3A2 | 9 | 9 | 19,4 | 64,9 | 27,64 | 599 | 0,68 | 1,01 |
| P49207 | 60S ribosomal | RPL34 | 3 | 3 | 16,2 | 13,3 | 28,45 | 117 | 0,68 | 1,36 |
| Q8N9Q2 | Protein SREK1 | SREK1IP1 | 4 | 4 | 22,6 | 18,2 | 27,69 | 155 | 0,65 | 1,03 |
| B4DEM9;Q9Y2 | Polymerase de | POLDIP2 | 2 | 2 | 7,1 | 39,9 | 23,98 | 350 | 0,64 | 0,62 |
| B4DLN1;P528 | 39S ribosomal | MRPL12 | 2 | 2 | 4,5 | 48,1 | 23,45 | 442 | 0,63 | 1,16 |
| H0Y2W2;Q9N | ATPase family | ATAD3A;ATAD | 7 | 7 | 12,6 | 64,2 | 27,74 | 572 | 0,62 | 1,24 |
| P62280;M0QZ | 40S ribosomal | RPS11 | 8 | 8 | 41,1 | 18,4 | 27,79 | 158 | 0,62 | 0,80 |
| P62913-2;P62 | 60S ribosomal | RPL11 | 5 | 5 | 31,1 | 20,1 | 27,04 | 177 | 0,61 | 0,86 |
| Q92930;H0YN | Ras-related pr | RAB8B | 3 | 2 | 18,8 | 23,6 | 24,10 | 207 | 0,61 | 0,76 |
| Q5T757;Q055 | Serine/arginin | SRSF11 | 2 | 2 | 4,2 | 48,5 | 25,33 | 424 | 0,60 | 0,56 |
| P49756 | RNA-binding p | RBM25 | 2 | 2 | 3 | 100,2 | 23,76 | 843 | 0,59 | 0,67 |
| G3XAC6;Q144 | RNA-binding p | RBM39 | 2 | 2 | 6,6 | 48,0 | 24,40 | 423 | 0,59 | 0,77 |
| M0R210;P622 | 40S ribosomal | RPS16;ZNF90 | 7 | 7 | 51,2 | 14,4 | 28,95 | 129 | 0,59 | 0,86 |
| E9PCT1;A9Z1 | Serine/arginin | SRRM1 | 2 | 2 | 2,6 | 93,4 | 23,49 | 820 | 0,57 | 0,51 |
| D6RBT3;O753 | NADH dehydr | NDUFS6 | 2 | 2 | 15,1 | 18,8 | 24,75 | 172 | 0,57 | 4,64 |
| F6VRR5;Q9BY | Polymerase de | POLDIP3 | 3 | 3 | 9,4 | 48,1 | 24,85 | 438 | 0,54 | 0,79 |
| Q7Z6Z7-2;Q7Z | E3 ubiquitin-p | HUWE1 | 3 | 3 | 0,9 | 480,2 | 24,68 | 4358 | 0,54 | 3,03 |
| P06576;H0YH | ATP synthase | ATP5B | 11 | 11 | 28,5 | 56,6 | 27,88 | 529 | 0,52 | 0,97 |
| H3BNX8;P206 | Cytochrome c | COX5A | 4 | 4 | 20,9 | 17,2 | 26,76 | 153 | 0,51 | 6,19 |
| P62987;M0R2 | Ubiquitin-60S | UBA52;UBB;R | 5 | 2 | 39,8 | 14,7 | 26,70 | 128 | 0,50 | 0,84 |
| A8MUH2;P18 | ATP synthase- | ATP5J | 2 | 2 | 26,1 | 14,0 | 25,25 | 119 | 0,49 | 1,08 |
| G3XAI9;Q5BK | Protein FAM1 | FAM133B;FAN | 2 | 2 | 17,1 | 12,4 | 24,99 | 105 | 0,49 | 0,56 |
| M0R0F0;P467 | 40S ribosomal | RPS5 | 5 | 5 | 34,5 | 22,4 | 26,58 | 200 | 0,49 | 0,75 |
| P63173;J3KT7 | 60S ribosomal | RPL38 | 4 | 4 | 50 | 8,2 | 27,59 | 70 | 0,48 | 0,85 |
| E9PPU1;P233 | 40S ribosomal | RPS3 | 5 | 5 | 46,8 | 17,4 | 27,43 | 158 | 0,48 | 0,99 |
| H0YLR3;H0YM | U2 small nucle | SNRPA1 | 7 | 7 | 60,7 | 9,5 | 29,99 | 89 | 0,48 | 0,67 |
| Q00325-2;Q0 | Phosphate car | SLC25A3 | 9 | 9 | 17,7 | 40,0 | 28,99 | 361 | 0,47 | 1,00 |
| P56385 | ATP synthase | ATP5I | 2 | 2 | 24,6 | 7,9 | 24,65 | 69 | 0,46 | 1,21 |
| P12236;Q9H0 | ADP/ATP tran | SLC25A6 | 11 | 6 | 31,2 | 32,9 | 27,55 | 298 | 0,41 | 1,27 |
| I3L1P8;Q0297 | Mitochondrial | SLC25A11 | 3 | 3 | 14,5 | 32,2 | 26,22 | 296 | 0,40 | 1,49 |
| Q96IX5 | Up-regulated | USMG5 | 2 | 2 | 27,6 | 6,5 | 24,57 | 58 | 0,40 | 0,64 |
| Q13523;H0YD | Serine/threon | PRPF4B | 3 | 3 | 5,1 | 117,0 | 25,32 | 1007 | 0,39 | 0,73 |
| M0R0Y3;Q9Y2 | RuvB-like 2 | RUVBL2 | 2 | 2 | 6,2 | 38,8 | 24,42 | 354 | 0,36 | 1,02 |
| Q13547;Q5TE | Histone deace | HDAC1 | 6 | 6 | 10,4 | 55,1 | 26,21 | 482 | 0,35 | 0,55 |
| P05141 | ADP/ATP tran | SLC25A5 | 8 | 3 | 24,5 | 32,9 | 28,07 | 298 | 0,31 | 0,85 |
| P10606 | Cytochrome c | COX5B | 3 | 3 | 28,7 | 13,7 | 26,88 | 129 | 0,29 | 4,86 |
| Q08AP5;C9J8 | Histone deace | HDAC10 | 1 | 1 | 14,6 | 11,2 | 24,11 | 96 | 0,17 | 0,80 |
